# Loss of cis-PTase function in the liver promotes a highly penetrant form of fatty liver disease that rapidly transitions to hepatocellular carcinoma

**DOI:** 10.1101/2023.11.13.566870

**Authors:** Abhishek K. Singh, Balkrishna Chaube, Kathryn M Citrin, Joseph Wayne Fowler, Sungwoon Lee, Jonatas Catarino, James Knight, Sarah Lowery, Sonal Shree, Nabil Boutagy, Inmaculada Ruz-Maldonado, Kathy Harry, Marya Shanabrough, Trenton Thomas Ross, Stacy Malaker, Yajaira Suárez, Carlos Fernández-Hernando, Kariona Grabinska, William C. Sessa

## Abstract

**Obesity-linked fatty liver is a significant risk factor for hepatocellular carcinoma (HCC)^1,2^; however, the molecular mechanisms underlying the transition from non-alcoholic fatty liver disease (NAFLD) to HCC remains unclear. The present study explores the role of the endoplasmic reticulum (ER)-associated protein NgBR, an essential component of the cis-prenyltransferases (cis-PTase) enzyme^3^, in chronic liver disease. Here we show that genetic depletion of NgBR in hepatocytes of mice (N-LKO) intensifies triacylglycerol (TAG) accumulation, inflammatory responses, ER/oxidative stress, and liver fibrosis, ultimately resulting in HCC development with 100% penetrance after four months on a high-fat diet. Comprehensive genomic and single cell transcriptomic atlas from affected livers provides a detailed molecular analysis of the transition from liver pathophysiology to HCC development. Importantly, pharmacological inhibition of diacylglycerol acyltransferase-2 (DGAT2), a key enzyme in hepatic TAG synthesis, abrogates diet-induced liver damage and HCC burden in N-LKO mice. Overall, our findings establish NgBR/cis-PTase as a critical suppressor of NAFLD-HCC conversion and suggests that DGAT2 inhibition may serve as a promising therapeutic approach to delay HCC formation in patients with advanced non-alcoholic steatohepatitis (NASH).**

## Introduction

Hepatocellular carcinoma (HCC) typically develops in patients with chronic liver disease resulting from viral (HBV and HCV) or non-viral (alcohol and nonalcoholic fatty liver disease (NAFLD)^4,5^. Recent epidemiological studies indicate an increase in HCC incidence in NAFLD patients, particularly in Western countries with elevated prevalence of obesity associated NAFLD^6^. NAFLD is characterized by excessive accumulation of triacylglycerol (TAG) in hepatocytes due to increased fatty acid (FA) uptake, biosynthesis, or imbalanced FA partitioning into storage, oxidative, and lipoprotein secretory pathways^7^. Sequential induction of nonalcoholic steatohepatitis (NASH) and hepatic fibrosis can occur in NAFLD, leading to HCC development.

Despite recent efforts to develop preclinical mouse models for NASH-HCC, including genetic, toxin, or dietary models, these models differ significantly in the cause of liver injury, penetrance, and length of onset of tumor development. Moreover, none of these models fully replicate human NAFLD-HCC pathology^8,9^. As a result, effective treatments for patients with NASH-HCC are currently unavailable, primarily due to a lack of precise understanding of the mechanisms underlying obesity-induced HCC in the context of NAFLD, and the absence of a reliable diet-induced HCC mouse model for hypothesis generation and therapeutic testing.

Previously, we have identified the highly conserved, ER-associated heteromeric protein complex responsible for cis-prenyltransferases (cis-PTase) activity, consisting of membrane associated subunits; nuclear undecaprenyl synthase (NUS1), also called NgBR and dehydrodolichol diphosphate synthase (Dhdds)^3,10,11^. This evolutionary conserved, heteromeric complex is essential for cis-PTase activity; the rate-limiting enzyme committed to the synthesis of dolichol phosphate (DolP), an obligate lipid carrier for protein glycosylation reactions including N-glycosylation, C-mannosylation, O-mannosylation, and GPI-anchor biosynthesis^3,12,13^. In recent years, the physiological significance of the NgBR/Dhdds complex has received considerable attention due to several in-depth genetic exome sequencing studies reporting various pathogenic mutations in both subunits and evidence that such loss of function mutations can cause diseases. Mutations within the genes of either subunit, such as the R290H loss of function mutation in NgBR or K42E mutation in Dhdds, can cause distinct clinical disorders ranging from the severe congenital disorder of glycosylation (CDG), retinitis pigmentosa, neurodegenerative diseases and epileptic encephalopathies^3,13-15^. Global deletion of NgBR in mice leads to early embryonic lethality (E 6.5) and conditional deletion in the endothelium causes lethality due to impairment of glycosylation of several critical proteins resulting in ER stress and impaired cell growth^3,10^. Collectively, these studies show that NgBR is essential for life via the critical role of dolichol-mediated protein glycosylation reactions that regulate the secretion, turnover and function of many proteins.

Defective or altered protein glycosylation is associated with the development of several cancers, including HCC^16-18^, which led us to investigate the role of NgBR in a mouse model of NAFLD. Surprisingly, deletion of NgBR in the liver (N-LKO) of mice fed a high-fat diet (HFD), but not a normal chow diet, promotes the hallmarks of fatty liver disease leading to HCC. This occurs with 100% penetrance after 16 weeks of HFD and the model exhibits relevant clinical features, including fatty liver secondary to impaired VLDL secretion, T-cell recruitment and fibrosis, ultimately resulting in carcinogenesis. Importantly, pharmacological inhibition of hepatic TAG synthesis in N-LKO mice prevents diet-induced metabolic alterations and HCC development, thus providing support for etiology of fat driven hepatic carcinogenesis.

## Results

### Liver-specific NgBR deficiency reduces VLDL production, promotes fatty liver disease and hepatocarcinogenesis

Previous studies have shown that liver-specific deletion of NgBR leads to insulin resistance and hepatic steatosis^19,20^. We were interested to understand this phenotype so we generated liver-specific knockout mice (N-LKO) **(Extended Data Fig. 1a,b)** and measured cis-PTase enzyme activity in hepatocytes. As anticipated, N-LKO mice displayed a marked reduction in cis-PTase activity, which confirmed the loss of enzymatic function **(Extended Data Fig. 1c)**. Next, hepatic fat content in N-LKO mice maintained on a chow diet (CD) for 24 weeks was measured and there was a fatty appearance, enhanced neutral lipid containing droplets and a two-fold increase in hepatic TAG content compared to wild-type (WT) littermates **(Figure 1a,b)**, supporting the idea that the loss NgBR in hepatocytes promotes steatosis.

**Fig. 1.**
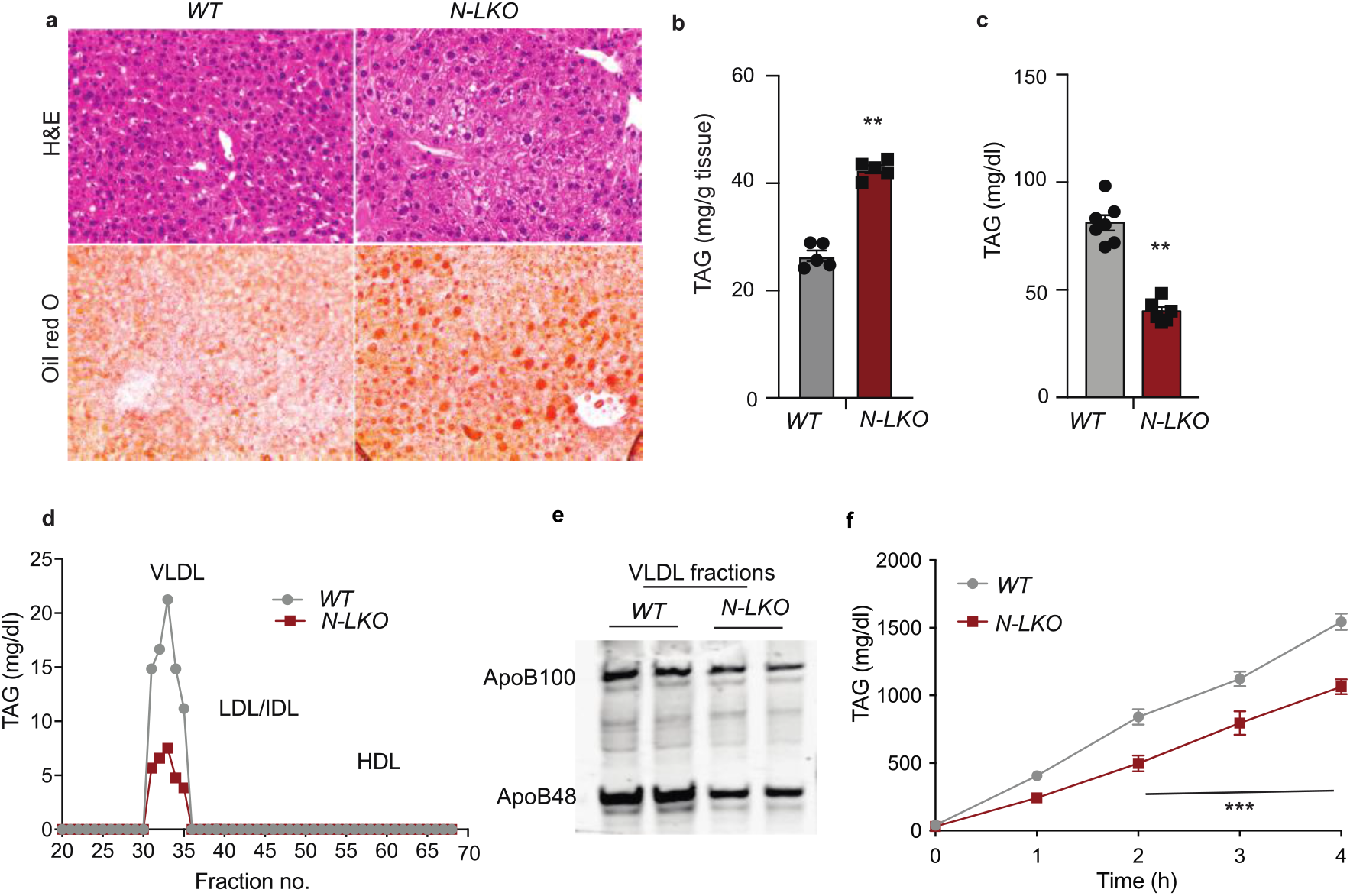
The loss of NgBR functions in the liver impairs hepatic VLDL-TAG secretion. **a,** Representative images of H&E-stained and oil red O-stained (ORO) liver sections, and **b,** hepatic TAG levels, both obtained from WT and N-LKO mice fed a chow diet (CD) for 6 months (n=4). **c,** Circulating TAG levels in overnight-fasted WT and N-LKO mice on a 6-month CD (n=6). **d,** TAG content of FPLC-fractionated lipoproteins from pooled plasma (n=5) of overnight-fasted WT and N-LKO mice fed a CD for 6 months. **e,** Representative western blot analyses of plasma ApoB100 and ApoB48 in the VLDL fractions from FPLC-fractionated lipoproteins (n=5). **f,** Plasma TAG levels in overnight fasted WT and N-LKO mice treated with LPL inhibitor 407 to block the lipolysis of circulating TAG-rich lipoprotein (n=5). Statistical significance: **P < 0.01 using Welch’s t-test **(b)** and ***P < 0.001 using two-way ANOVA with Sidak’s multiple comparisons test **(f)**.

To elucidate the mechanism by which hepatic NgBR deficiency promotes liver fat accumulation, we assessed circulating TAG levels in N-LKO mice maintained on a CD for 24 weeks or on a high-fat diet (HFD) for 16 weeks. Interestingly, both groups demonstrated significantly decreased plasma TAG levels under fasting conditions **(Figure 1c and Extended Data Fig. 2a)**. FPLC analysis revealed reduced TAG levels in VLDL fractions and decreased apolipoprotein ApoB100 and ApoB48 levels in isolated VLDL fractions for N-LKO mice compared to WT counterparts **Figure 1d,e and Extended Data Fig. 2b)**. To determine the cause of reduced plasma VLDL-TAG in N-LKO mice under physiological and pathological conditions during fasting, we assessed VLDL secretion by inhibiting lipoprotein lipase in N-LKO mice. N-LKO mice exhibited a substantially lower rate of hepatic VLDL-TAG production **(Figure 1f)** compared to controls suggesting that the decreased plasma TAG levels in N-LKO mice are a consequence of impaired hepatic VLDL secretion, which may subsequently contribute to increased fat accumulation in the liver and the onset of steatosis.

Next, systemic metabolic function in N-LKO and control mice subjected to a high-fat diet (HFD) for 16 weeks was assessed. Interestingly, N-LKO mice displayed significant decreases in body weight, food consumption, and water intake, while no differences were observed in energy expenditure, CO2 production, O2 consumption, respiratory exchange ratio, or locomotor activity **(Extended Data Fig. 3a-e and Extended Data Fig. 4a-f)**. These findings point to a hypophagic response in these animals, which may have implications for liver health.

Upon sacrifice, remarkably, both male and female N-LKO mice exhibited a 100% incidence of multiple liver tumor nodules when fed either HFD or high-fat cholesterol Western diet (WD-western type diet) for 16 weeks **(Figure 2a & Extended Data Fig. 5a)**, indicating a potential connection between the hypophagic state and the development of liver tumors. This phenomenon was observed in offspring from at least three independent cohorts of mice but was not detected in N-LKO mice fed a regular CD, even when sacrificed approximately 20 months after birth **(Extended Data Fig. 5b)**. Histological analysis revealed that liver tumors consisted of both the fatty transformation of cells as well as solid growth of tumors with distinct margins **(Figure 2a)**. Next, we examined hepatic cell proliferation in tumors by Ki-67 staining. Compared to the adjacent non-tumorous liver, the tumor cells showed markedly increased levels of Ki-67 **(Figure 2b)**; resembling NAFLD initiated HCC. Consistent with the development of HCC, alpha fetoprotein (AFP), a plasma biomarker for proliferating hepatocytes and HCC, was significantly elevated in high-fat fed N-LKO mice **(Figure 2c)**. Assessment of blood cell counts after HFD for 16 weeks revealed that circulating WBC/lymphocytes were significantly increased in N-LKO mice **(Figure 2d)** and the proportion of other circulating cells were similar in both groups of mice **(Figure 2d)**. Profiling of lymphocyte subsets showed that CD8+T cells were increased amongst the lymphoid population in N-LKO mice fed with HFD **(Figure 2e)**. To examine if NgBR levels are impacted by NAFL/NASH transition, human transcriptomic datasets were interrogated. As seen in **(Figure 2f)**, hepatic NgBR transcript levels are progressively diminished in patients with NAFL and NASH across different fibrosis stages (F0-F1, F3, and F4) compared to healthy individuals **(Figure 2f)**. The consistency between the murine model and human data strengthens the hypothesis that suppression of NgBR in late-stage NASH, a pre-HCC condition, may potentially contribute to liver cancer development.

**Fig. 2.**
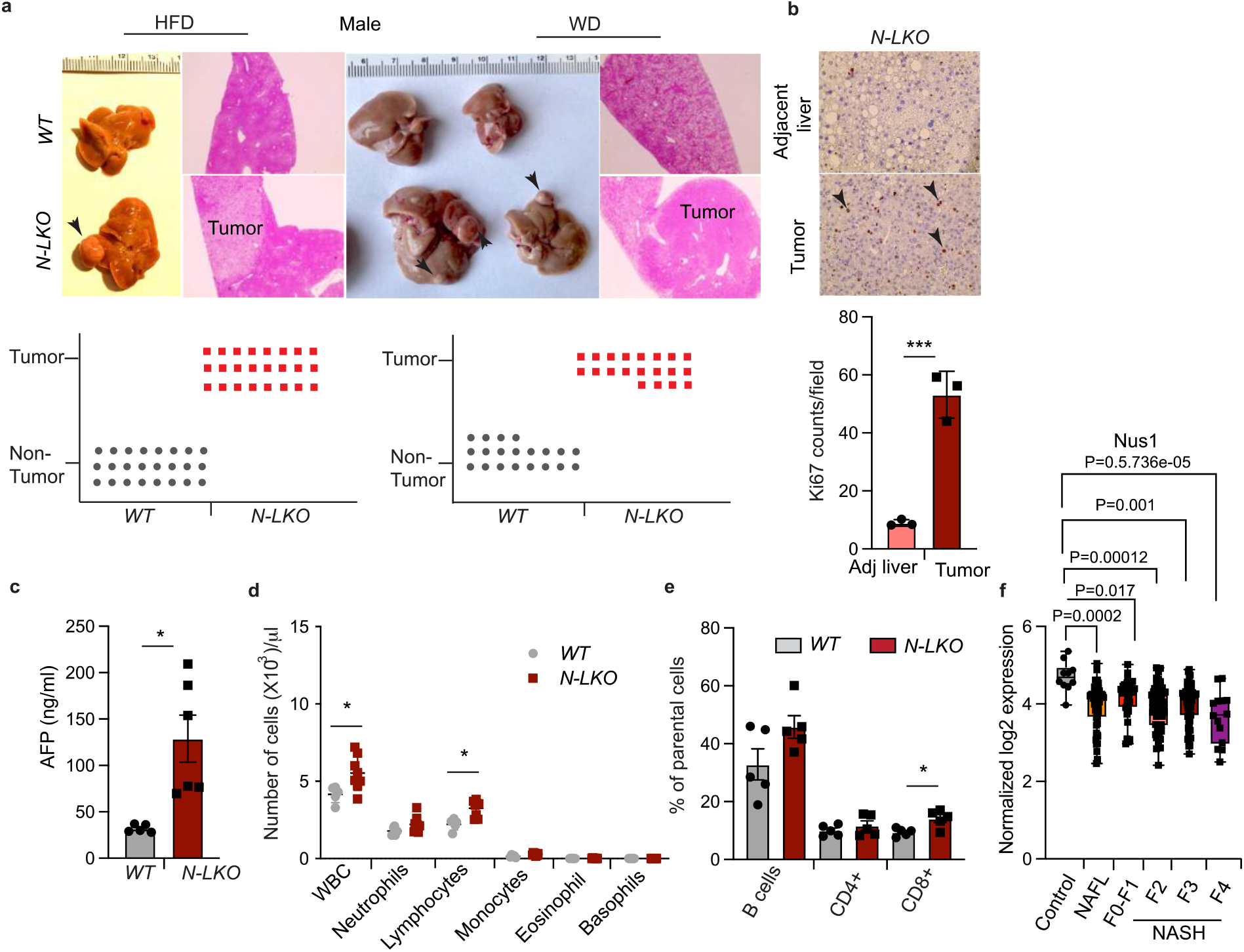
Loss of hepatic NgBR drives HCC development in diet-induced obesity and correlates with NAFLD-NASH progression in human liver. **a,** Representative images of livers from wild-type (WT) and liver-specific NgBR-deficient (N-LKO) male mice fed a high-fat diet (HFD) or western-type diet (WD) for 16 weeks, with arrows indicating HCC. Histological analysis of liver and tumor sections stained with H&E are shown in the right panel, and the graph summarizes the incidence of HCC (lower panel) in N-LKO and WT mice on HFD (n=24) and WD (n=16). The scale bar represents 200 μm and the icons indicate individual mice. **b,** IHC analysis of the proliferation marker Ki-67 in tumor and tumor-adjacent liver of N-LKO mice kept on WD for 16 weeks, with quantification of Ki-67/field (n=3) shown in the lower panel. Scale bar, 50 μm; **c,** Circulating AFP levels in WT and N-LKO mice fed HFD (n=5,6). **d,** Peripheral blood cell counts from WT and N-LKO mice fed HFD for 16 weeks, measured using a hemavet hematology analyzer (n=5). **e,** Flow cytometry analysis of circulating B and T cells (n=5). **f,** The log2 normalized mRNA expression levels of NgBR (Nus1) in human liver samples with non-alcoholic fatty liver (NAFL) and varying stages of non-alcoholic steatohepatitis (NASH) with different degrees of fibrosis. The expression levels were obtained through RNA sequencing of 15 healthy liver samples, 30 steatosis samples, and 20 NASH samples at stages F0-F4. The analysis was adjusted for P values. Each data point represents a biological replicate. All data are represented as mean ± SEM, and two-sided *P < 0.05; ***P < 0.001, comparing N-LKO with WT mice using an unpaired Welch’s t-test (**b-e**).

To independently validate the concept that a loss of cis-PTase activity functions as a driver of advanced liver disease, we took two additional approaches to reduce cis-PTase activity in mice. Previously, it was shown that a NgBR R290H mutation in humans is a loss off function mutation by virtue of this amino acid substitution impairing substrate binding and enzymatic function of the NgBR/DHDDS complex, causing a severe congenital disorder of glycosylation. Thus, we generated NgBR orthologous R294H mutant knock-in mice **(Extended Data Fig. 6a)**. Homozygous R294H NgBR mice die ∼4-12 weeks after birth while heterozygous R294H NgBR mice were born with normal frequencies **(Extended Data Fig. 6b)**. Interestingly, HCC developed in heterozygous NgBR R294H mutant mice (incidence∼20%) fed with HFD for 16 weeks **(Extended Data Fig. 6c-e)**. Secondly, we generated mice lacking hepatic Dhdds (D-LKO) **(Extended Data Fig. 7a-b)**, an obligate subunit of enzyme cis-PTase that heterodimerizes with NgBR. Feed D-LKO mice a HFD for 16 weeks also promoted HCC with a frequency of ∼40% **(Extended Data Fig. 7c)**, suggesting a critical role of cis-PTase activity in preventing hepatocyte transformation when exposed to a lipid-rich diet. Collectively, our data demonstrate that hepatic loss of function of the heteromeric NgBR/DHDDS cis-PTase complex in three distinct genetic models directly contributes to liver carcinogenesis after a short duration of feeding mice a HFD.

### Hepatic ablation of NgBR induces genomic instability

Having shown a role for NgBR regulating hepatic responses to diet-induced liver carcinoma, we wanted to discern the mechanistic underpinnings of this observation. Due to the complicated link between fatty liver disease and HCC development, the molecular drivers involved in hepatocarcinogenesis are inadequately understood. To begin to understand the evolutionary forces and molecular basis driving NAFLD-HCC, we performed whole-genome sequencing of isolated tumor tissue and adjacent regions of the liver of N-LKO mice fed a WD for 16 weeks. Surprisingly, mutations were found in both the tumor and adjacent liver specimens, compared to DNA sequenced from the liver of isogenic, control C57BL6 mice and a higher mutation burden was found in tumor versus tumor-adjacent DNA **(Extended Data Fig. 8a)**. Furthermore, we analyzed all 96 possible base substitutions in the non-tumor adjacent liver and tumor based on the trinucleotide framework. Among these single nucleotide variants (SNV) mutations, C to T transitions were the most frequent SNV, followed by T to C conversions **(Extended Data Fig. 8b)**. This mutation profile is consistent with the previously identified signature for human and mouse liver cancer^21,22^. In addition, there were mutations in eight protein-coding regions of genes in the non-tumor adjacent liver **(Extended Data Fig. 8c)** and an excess of missense mutations was observed in mucin 4 (*muc4*) **(Extended data Fig. 8c)**. *Muc4* is one of the most repeatedly mutated genes in HCC patients and is a major transmembrane mucin family member that is extensively glycosylated^23,24^. Western blotting for of MUC4 from livers of N-LKO as compared to WT mice documented its hypoglycosylation by accumulation of the lower apparent molecular weight species **(Extended data Fig. 8d)**. Thus, the loss of hepatic NgBR triggers several mutations in *muc4* gene and impairs glycosylation of MUC4 protein. Additionally, a frame-insertion occurs in the *blm gene*, which encodes the Bloom syndrome protein related to RecQ helicase, a protein involved in maintaining the DNA topology during the DNA replication and repair processes **(Extended Data Fig. 8c)**. Mutations detected in *muc4 and blm* are consistent with previous reports in human HCC^23,25,26^. To our knowledge, the other six hepatic genes mutated **(Extended Data Fig. 8c)** with missense, in-frame insertion, and frameshift mutations have not been previously reported in human HCC. These results indicate that mice lacking NgBR in the liver promotes genomic instability to drive the diet-induced HCC development.

### Hepatic NgBR deficiency upregulates gene expression profiles associated with HCC

To assess global pattens of gene expression, we performed unbiased RNA-seq analysis on RNA isolated from livers of N-LKO and WT mice fed a HFD for 16 weeks. Multidimensional scaling (MDS) plot analysis of HFD-fed mice reveals a clear separation of gene expression patterns based on genotype **(Extended Data Fig. 9a)**. In N-LKO mice, there were 697 genes whose expression was differentially regulated (DE), among which 407 genes were significantly upregulated and 290 were down-regulated **(Extended data Fig. 9a)**. We then used the Kyoto Encyclopedia of Genes (KEGG) pathway analysis to evaluate the comparative function of these dysregulated genes. We identified numerous genes involved in promoting canonical pathways such as fatty liver formation, ROS production, DNA damage, and cancer development that were upregulated in N-LKO compared to WT mice **(Extended Data Fig. 9b)**. Several genes of these pathways are thought to play a critical role in HCC development and may contribute to impaired liver functions and tumor formation in N-LKO mice. Similar to hepatic transcriptome analysis, we have also performed unbiased RNA seq analysis from tumors of WD fed N-LKO mice compared to WD fed livers of WT mice. Heatmap analysis of the top 20 altered genes in tumors of NLKO mice confirmed upregulation of AFP and FABP5 or downregulation of PCSK9 cancer or cancer-associated genes **(Extended Data Fig. 9c)** as markers responsible for promoting tumor development.

To investigate liver cell heterogeneity and dynamic alterations during NAFL-HCC pathogenesis, single-cell RNA sequencing on parenchymal (PCs) and non-parenchymal cells (NPCs) or tumor cells isolated from the livers of WT or N-LKO mice fed a WD for 16 weeks was performed. Data analysis identified 33 discrete cell populations clustered into 11 unique cell types, including hepatocytes, hepatic stellate cells (HSCs), cholangiocytes, endothelial cells, macrophages, dendritic cells, Kupffer cells, T cells, and B cells **(Figure 3a and Extended Data Fig. 10a)**. By delineating component cell analysis, a distinctive transcriptional signature of PCs and NPCs in N-LKO compared to WT mice identified nine distinct hepatocyte clusters comprising Hep1-Hep9, with two unique hepatocyte clusters observed only in N-LKO mice **(Extended Data Fig. 10a-c)**. Furthermore, the Hep4 population in N-LKO mice was enriched for HCC, as confirmed by high expression of cytokeratin genes such as Krt8, Krt18, Spp1 and Epcam **(Figure 3a)**. Gene set enrichment analysis revealed that the hepatocytes in N-LKO mice had decreased lipoprotein metabolism, mitochondria function and oxidative stress response compared to WT mice **(Figure 3b-c, Extended Data Fig. 10d and Extended Data Fig. 11a-d)**. Additionally, there was marked upregulation of genes associated with ER stress **(Extended Data Fig. 11e)**, consistent with loss of NgBR as previously described^10^. We also identified a decrease in growth suppressors and cell cycle checkpoints enriched genes, such as APC, p53 and PTEN **(Extended Data Fig. 12a-f)**. and enhanced expression of genes related to oncogenic signaling pathways, including receptor tyrosine kinase signaling (EGFr, FGFr2, INSr, NRP and BRAF), Rac/Rho GTPases, VEGF signaling, and extracellular matrix remodeling **(Extended Data Fig. 13a-h)**. These findings collectively suggest that the observed changes in gene expression may contribute to the development of cancer in N-LKO mice.

**Fig. 3.**
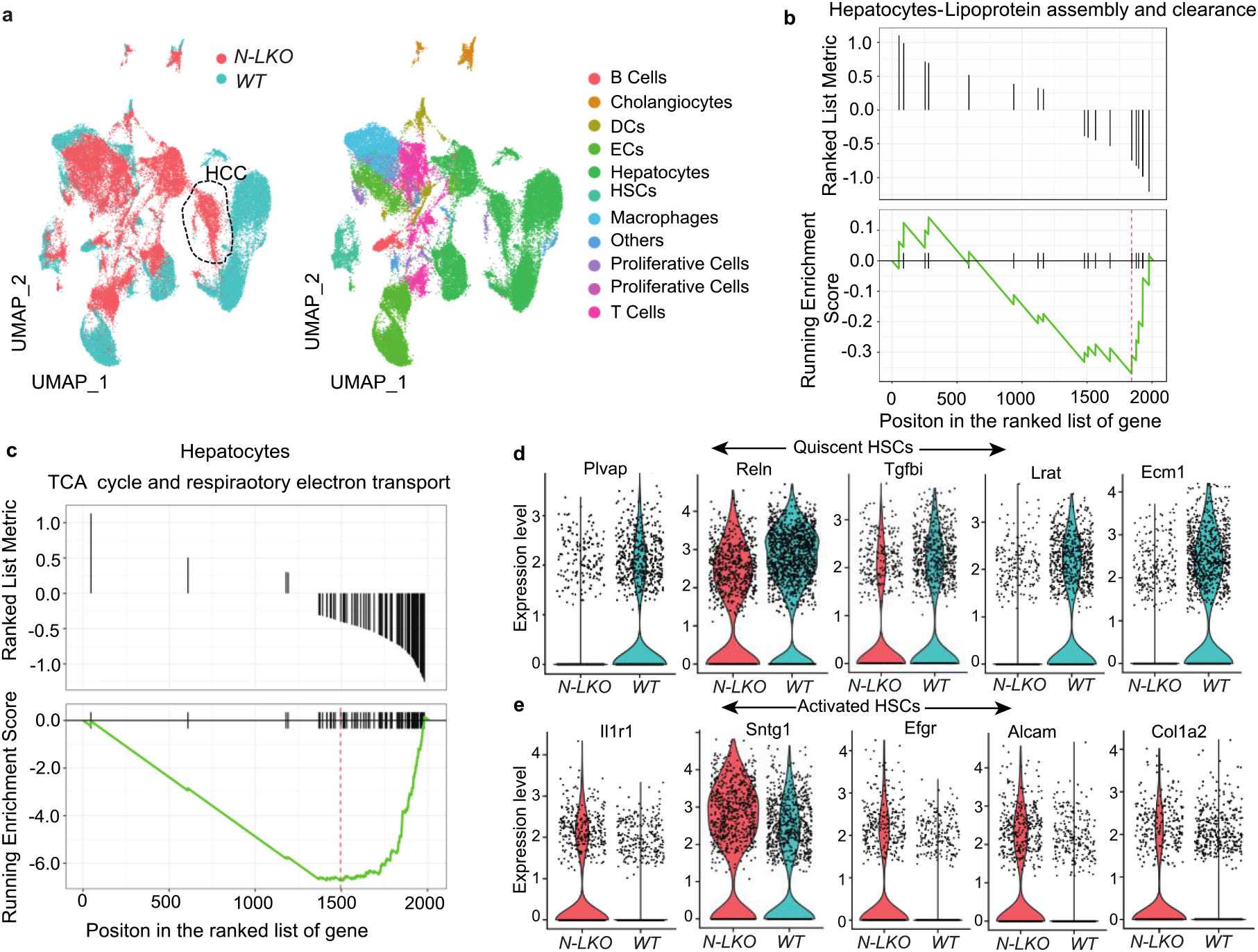
Liver-specific NgBR depletion induces gene expression patterns associated with NAFLD and HCC in hepatic cells. **a,** Single-cell RNA sequencing of the livers isolated from WT and liver-specific NgBR knockout (N-LKO) mice using the 10x Genomics Chromium platform. Uniform manifold approximation and projection (UMAP) visualize clustering of liver single-cell transcriptomes (7000 cells from WT and 7000 cells from N-LKO mice fed WD). Color annotating cell type or genotypes, circle around cluster represent cancer cells. **b-c,** Enrichment Score curves Gene set enrichment analysis (GSEA) plots of significant differentially expressed genes between WT and N-LKO hepatocytes. The peak in the plot shows the downregulation of the gene sets associated with Lipoprotein packaging and secretion and mitochondrial oxidative function. **d**-**e,** The violin plots compare the expression levels of hepatic stellate cell (HSC) quiescent marker genes (upper panel) and activation marker genes (lower panel) that are downregulated and upregulated, respectively, in N-LKO mice relative to WT mice. The plots illustrate a significant shift in the phenotype of hepatic stellate cells in N-LKO mice, as demonstrated by the altered expression of quiescent and activation markers.

We also investigated other important non-parenchymal cells in the liver, particularly T cells and HSC in both WT and N-LKO mice. Analysis showed that there is a marked difference in gene expression among all four T cell subclusters in N-LKO mice compared to WT mice **(Extended Data Fig. 14a-c)**. In addition, a notable increase in T cell markers across all clusters exhibiting reduced enrichment of genes linked to fatty acid metabolism, mitochondrial function, and oxidative stress response were found in N-LKO mice **(Extended Data Fig. 14d-e)**. This rise was accompanied by an increase in T cells expressing markers of stress and exhaustion. **(Extended Data Fig. 14f-g)**. suggesting chronic inflammation, a hallmark of NASH. Furthermore, analysis of HSCs identified a subcluster **(Extended Data Fig. 15a,b)** that had reduced expression of HSC quiescent markers, lipid metabolism-related genes and oxidative stress response genes in N-LKO mice compared to the control group **(Figure 3d and Extended Data Fig. 15c-e)**. In contrast, this subcluster of HSCs exhibited higher expression of markers associated with HSC activation and genes enriched in extracellular matrix remodeling **(Figure 3e and Extended Data Fig. 15f-h)**, indicating that N-LKO mice may be at a higher risk for liver fibrosis. Collectively, these changes in impaired liver lipoprotein metabolism, oxidative stress, inflammation, and fibrosis may contribute to liver dysfunction and HCC formation observed in these mice.

### Hepatic NgBR suppression promotes TAG accumulation, inflammation, and fibrosis in the liver

Utilizing single-cell RNA sequencing analysis, we identified that NgBR deficiency in mice liver led to impaired lipid metabolism pathways, augmented immune cell infiltration markers, and upregulated expression of genes related to ER and oxidative stress as well as fibrosis. These are important characteristics of NAFL-derived HCC^27^. To further explore the functional consequences of these observations, we characterized the fatty liver phenotype and specific immune cell subsets present in the liver of N-LKO mice subjected to a 16-week high-fat diet (HFD) regimen. Consistent with previous reports^19^, N-LKO mice fed a HFD displayed extensive hepatic TAG accumulation **(Figure 4a,b)**. HFD-fed N-LKO mice exhibit significantly increased hepatic infiltration of lymphocytes such as CD8+ T and NK cells, and myelocytes, including macrophages, dendritic cells (DCs), and neutrophils, but no alteration in B cells, CD4+T cells, and monocytes **(Figure 4c,d and Extended Data Fig. 16a-f)**. Previous studies have reported that inflammation found in fatty liver disease is accompanied by the hepatic infiltration of resident CD8 + T cells that express programmed death protein (PD-1)^28^, thus, suppressing the anti-cancer action of cytotoxic CD8+ T cells and driving the development of HCC. Indeed, CD8+ T cells that express elevated levels of PD-1 are found in the liver of HFD-fed N-LKO mice **(Extended Data Fig. 16g)**. Furthermore, single-cell RNA sequencing analysis in hepatocytes demonstrated a significant elevation in acute phase response genes, including Saa1, Mt1, Mt2, Lcn2, and Orm2, which arise from inflammation in N-LKO mice compared to WT mice fed WD **(Figure 4e)**, consistent with previous reports^29,30^. Concomitantly, N-LKO mice fed on HFD for 16 weeks had increased liver damage, as shown by increased hepatic collagen deposition and fibrosis (Sirius red staining), augmented expression of fibrotic genes and elevated levels of liver-damaging enzyme ALT and AST in the plasma, markers of liver damage resembling the pattern in human NASH **(Figure 4f-i)**. Excessive hepatic lipid accumulation increases inflammation, fibrosis, and oxidative DNA damage caused by elevated ROS production, lipid peroxidation, and ER stress, well-established drivers of liver damage^31-33^. N-LKO mice fed with HFD show significantly increased hepatic ROS production, MDA content (final oxidative product of polyunsaturated FA peroxidation), and key ER stress marker such as ATF4 and Bip1 **(Extended Data Fig. 17a-c)**. These results imply that hepatic suppression of NgBR contributes to the aggravation of NAFLD, NASH, and fibrosis, which may account for the HCC development.

**Fig. 4.**
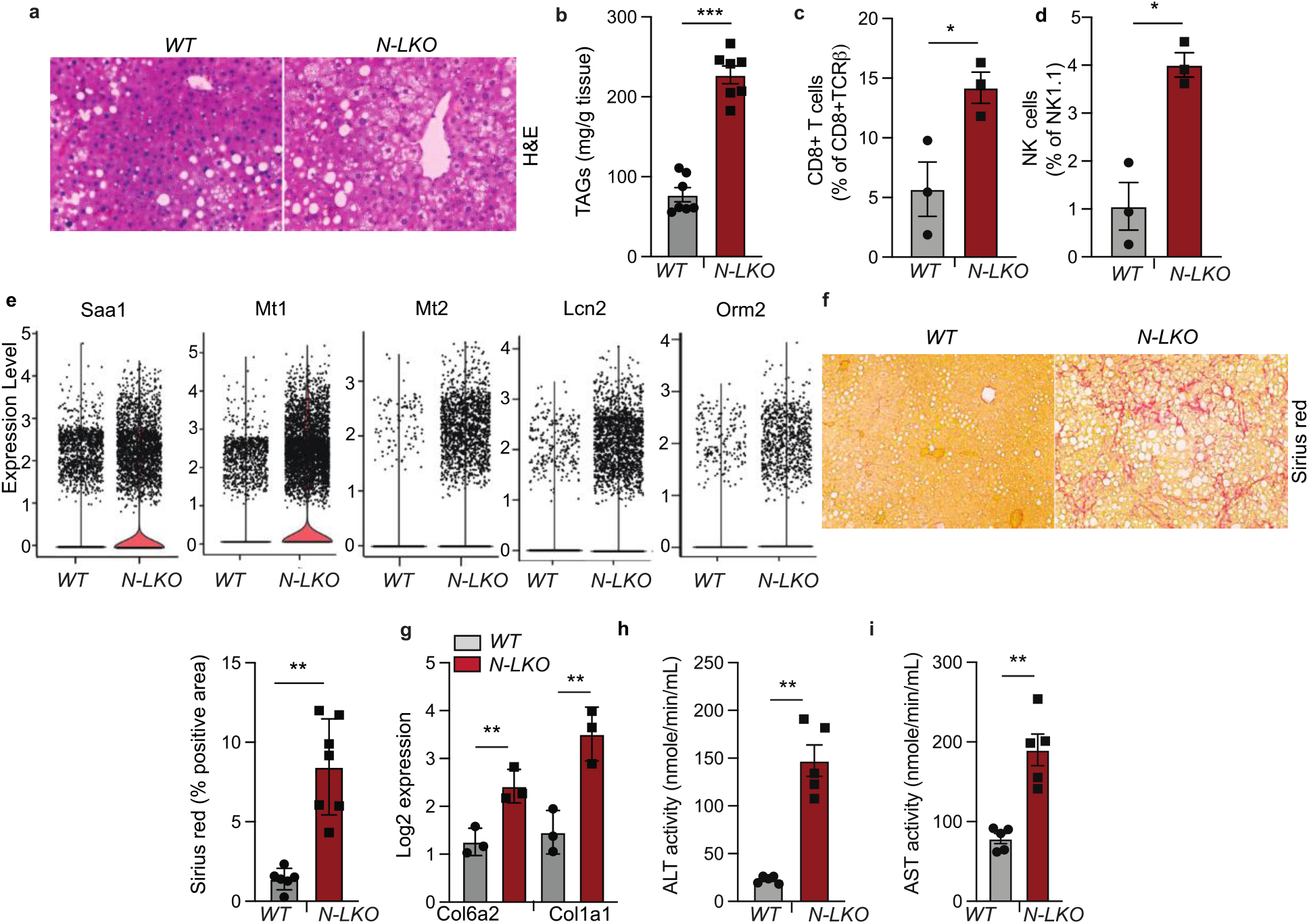
Hepatic NgBR deficiency exacerbates NASH-fibrosis phenotype in response to overnutrition. **a,** Representative images of liver sections stained with H&E, and **b,** hepatic TAG levels in WT and N-LKO mice fed a HFD for 16 weeks (n=7). Scale bar, 50 μm. (**c-d**) Liver lymphoid cells including CD8+ T cells and NK cells (n=3) isolated from WT and N-LKO mice fed an HFD for 16 weeks assessed by flow cytometry. **e,** The violin plots illustrate the expression of acute phage response genes is elevated in the hepatocytes of N-LKO compared WT mice that were fed a WD. **f,** Representative images and quantification of Sirius red staining in the liver sections from WT and N-LKO mice fed HFD for 16 weeks (n=7). Scale bar, 100 μm **g,** Log transformed mRNA expression of Col6a2 and Col1a1 from liver of WT and N-LKO mice fed an HFD for 16 weeks (n=3). **h-i,** Plasma ALT and AST levels in WT and N-LKO mice fed an HFD (n=5). All data are represented as mean ± SEM. Two-sided \**P* < 0.05; \*\**P* < 0.01; \*\*\**P* < 0.001, comparing N-LKO– with WT mice using an unpaired Welch’s t-test.

### DGAT-2 inhibition blocks obesity-induced HCC development in N-LKO mice

Based on in vivo data, it is evident that mice lacking NgBR in hepatocytes fed a HFD promotes lipid overload, leading to cellular reprogramming that may favor DNA damage and the development of HCC. To directly examine if attenuation of liver fat accumulation impacts the progression to HCC, N-LKO mice were treated with a small molecule inhibitor (PF-06424439) of diacylglycerol acyl transferase-2 (DGAT2). In rodents and primates, DGAT2 catalyzes the final rate-limiting reaction of TAG biosynthesis by incorporating the last fatty acyl group to diacylglycerides^34^. PF-06424439 inhibits DGAT2 and substantially reduces circulating TAG levels and hepatic fat in rodents with NAFLD^35^. Thus, 8-week old N-LKO mice were fed a WD formulated with or without PF-06424439 for 16 weeks. Administration of PF-06424439 in N-LKO mice (N-LKO+DGI) markedly reduced hepatic TAG contents and fibrosis **(Figure 5a,b)**, consistent with previous reports in rodents and humans^34,35^. PF-06424439 treated N-LKO mice were also protected against WD-induced liver damage, as shown by lower circulating levels of liver enzyme ALT and AST **(Figure 5c,d)**. We next determined whether lowering TAG levels in the liver by DGAT2 inhibition attenuates lipid-mediated deleterious effects in NLKO mice fed with WD. PF-06424439 treatment in NLKO mice fed with WD significantly reduced hepatic ROS production, MDA content, and ER stress marker such as ATF4 **(Figure 5e,f,g)** and completely abolished HCC development and circulating AFP levels **(Figure 5h,i)**.

**Fig. 5.**
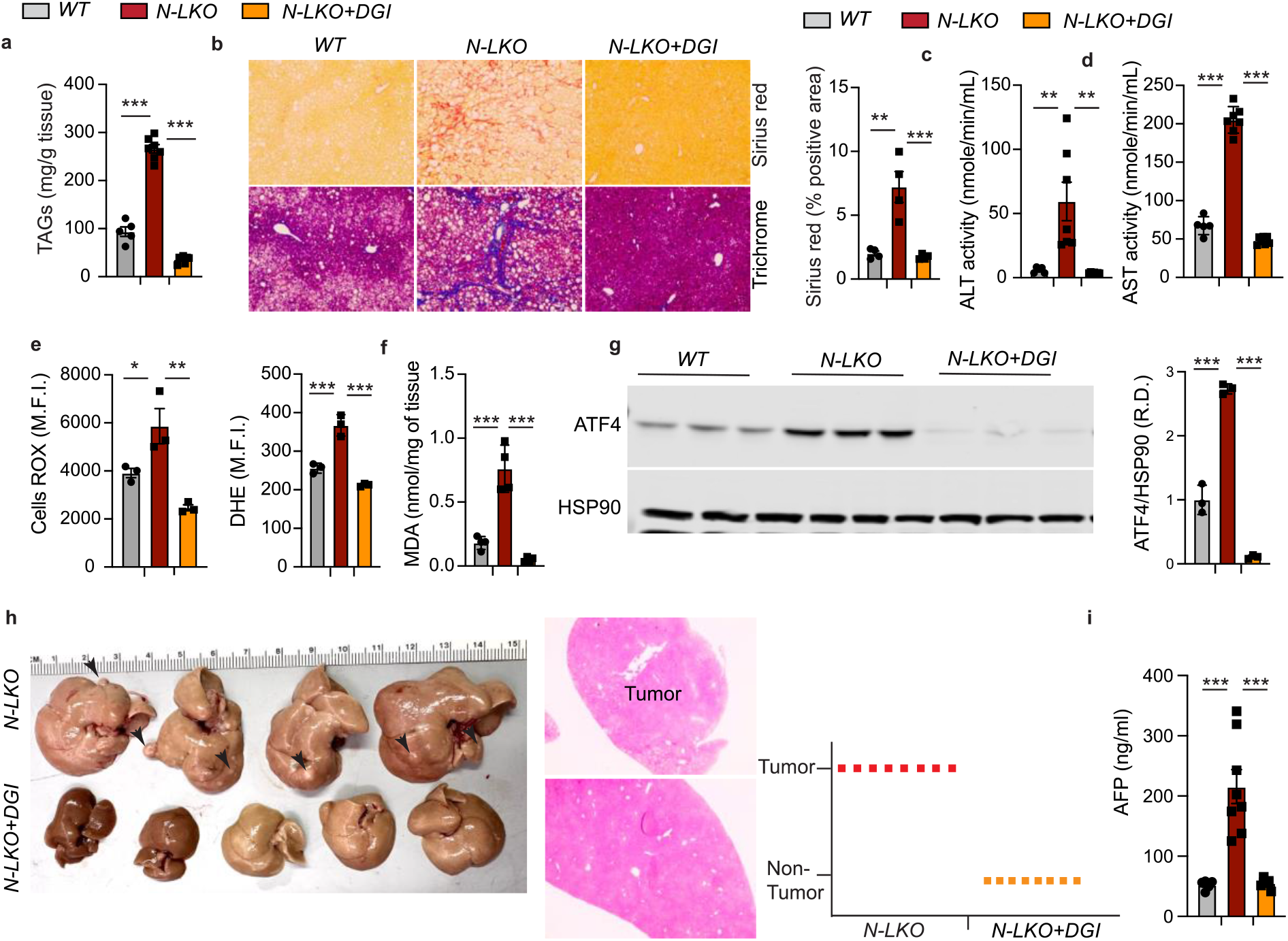
DGAT2 inhibitor treatment prevents the diet-induced NAFL-HCC pathogenesis. This study involved three groups of mice: WT mice receiving Western diet (WD) for 16 weeks and N-LKO mice of the same age group, with N-LKO divided into two subgroups - one receiving a WD only and the other receiving formulated WD with a DGAT2 inhibitor (designated as N-LKO+DGI) for 16 weeks. After 16 weeks, the following parameters were measured. **a,** Hepatic TAG levels in all three groups of mice (n=5). **b,** Representative images and quantification of Sirius red staining (Upper and right panel) and Trichome staining (lower panel) in liver sections from all three groups of mice (n=5). Scale bar, 100 μm. **c-d**, Serum ALT and AST levels in all three groups of mice (WT n=5, N-LKO n=6, N-LKO+DGI n=7). **e,** Analysis of cellular ROS in primary hepatocytes isolated from all three groups of mice (n=3). **f,** Membrane lipid peroxidation determined through MDA assay in the liver of all three groups of mice (n=4). **g,** Representative western blot and densitometric analysis of a key ER stress response protein ATF4 and housekeeping standard HSP90 in all three groups of mice (n=3). **h,** Representative photographs of livers isolated from N-LKO and N-LKO+DGI mice, with histological analysis of liver and tumor sections stained with H&E, and a graph summarizing the occurrence of tumors in both groups (n=8). Scale bar, 200 μm. **i,** Plasma AFP levels in all three groups of mice (WT n=5, N-LKO n=8, N-LKO+DGI n=8). The bar with dot plots represents the mean ± SEM, with each point representing a biological sample. Statistical analysis was performed using one-way ANOVA with Tukey’s multiple comparisons test (*P < 0.05; **P < 0.01; ***P < 0.001).

Taken together, these results show that the loss of NgBR and a reduction is cis-PTase activity in the liver promotes elevated hepatic TAG levels and many of the hallmarks of NAFLD driving genomic instability and liver cancer rapidly and in a remarkably highly penetrant manner. Pharmacological inhibition of DGAT2 restores normal physiological liver lipid homeostasis, resulting in comprehensive prevention of HCC development in NLKO mice. These findings support and give credence to the possibility that reducing liver fat accumulation may impact the transition from NAFLD to HCC.

## Discussion

The development of HCC in patients with fatty liver is a major concern, increasing in frequency in patients with late-stage NASH. However, the mechanisms underlying this progression are not yet fully understood and difficult to model experimentally. To shed light on this issue, we found that liver-specific deletion of NgBR aggravated HFD driven inflammation, oxidative and ER stress, and fibrosis, leading to the development of liver cancer with 100% incidence, thus providing a robust model of NAFL-HCC transition. Since these phenotypes did not occur in NgBR deficient mice on a normal chow diet, and HFD driven HCC was attenuated by pharmacologic inhibition of DGAT2, the data imply that cis-PTase activity plays a critical role in suppressing NAFLD-linked HCC by maintaining liver triacylglycerol (TAG) homeostasis.

Mechanistically, since hepatic NgBR deficiency reduces glycosylation of many intracellular and secreted proteins leading to the unfolded protein response, a synergy of multiple hits, triggered by HFD may explain the highly penetrant phenotype. In the case of lipid overload, the reduced secretion of VLDL can lead to abnormal lipid accumulation in the liver, triggering the production of reactive oxygen species and lipid peroxidation, causing oxidative stress. Furthermore, suppressing NgBR can lead to reduced protein glycosylation in the ER, which impairs protein folding and triggers ER stress. The combined effects of increased oxidative, and ER stress can lead to DNA damage and mutations, ultimately resulting in the development of liver cancer. Indeed, the hepatic loss of NgBR leads to increased liver TAG accumulation as the primary cause of liver pathology and HCC, since blocking TAG synthesis with a DGAT2 inhibitor prevented HCC development and largely dampened the severity of lipotoxic-induced liver damage and fibrosis, demonstrating that TAG accumulation is the primary driver of HCC development in this model.

In recent years, various diet-induced HCC mouse models have been reported, including choline deficient HFD (CDHFD)^36^, high fat/sugar diets that induced spontaneous NASH-HCC in male B6/129 mice^37^, chemical-induced models^38^, and hepatocyte-specific deletion or overexpression models of PTEN^39,40^ or urokinase plasminogen activator (uPA)^41^. However, these models have limitations, such as low penetrance of liver tumor incidence, liver toxins, prolonged duration, and mixed genetic backgrounds. In addition, some models, such as DEN and CCl4, cause liver damage in a uniform manner^2,42^ that may not replicate the complex and dynamic progression of NAFLD-NASH. To address these limitations, our study provides a reliable and efficient diet-induced HCC mouse model that overcomes the limitations of previous models. This model offers potential targets for therapeutic intervention and provides a valuable tool for gaining insights into the mechanisms underlying HCC development.

Overall, our findings provide significant insights into the pathogenesis of NAFLD-NASH-HCC and suggest potential therapeutic targets for this disease. The reliable diet-induced HCC mouse model presented in this study could facilitate better understanding of obesity-induced HCC prevention and provide a valuable tool for advancing research in this field.

## Methods

### Animal studies

Generation of liver-specific NgBR or Dhdds-deficient mice. Mice bearing a loxP-flanked NgBR allele (*NgBR^loxP/loxP^* mice) or Dhdds allele (*Dhdds^loxP/loxP^*mice) were generated as described previously^3,43^. Liver-specific NgBR or Dhdds–knockout mice (N-LKO or D-HKO) were generated by breeding albumin-Cre; *NgBR^loxP/+^* mice or albumin-Cre; *Dhdds^loxP/+^* with *NgBR^loxP/+^* mice or *Dhdds^loxP/+^* mice respectively. All mouse strains were in the C57BL6 genetic background. *N-LKO or Dhdds* mice were confirmed for NgBR KO or Dhdds KO in the liver by PCR using Cre primers and primers flanking the 5′ homology arm of the NgBR gene or Dhdds gene and LoxP sites from the tail-extracted DNA. Heterozygous NgBR R294H mutant mice were created on C57BL6 background by the knock-in (KI) technology at Yale. PCR primers were designed that covered the targeted region (Forward primer, 5′-TCTAGGCTCTGTCACCCGCA-3′ and Reverse primer 5′-TCTAGGCTCTGTCACCCGCA-3′) amplifying a 269 bp fragment of DNA in both WT and NgBR R294H mice. The knock-in sequence was confirmed by restriction enzyme digestion with BStN1, that cleaves the knock-in allele.

All experimental mice were housed in a barrier animal facility with a constant temperature and humidity in a 12-hour dark and light cycle while water and food were provided *ad labium*. All mice (n=3-5 per cage) were fed with a standard chow diet (CD) for 8 weeks after that switched to an HFD (60% calories from fat; Research Diets D12492) or high fat high cholesterol, western diet (WD) containing 1.25% cholesterol (D12108, Research Diets) for 4 months to induce hepatocellular carcinoma (HCC).). Mice used in all experiments were sex- and age-matched and kept in ventilated cages in a pathogen-free facility. All the described experiments were approved by the institutional animal care use committee of Yale University School of Medicine.

### Metabolic cage analysis

For metabolic cage studies, mice were exposed to comprehensive lab animal monitoring system (CLAMS; Columbus Instruments) to profile food consumption, water intake, oxygen consumption (VO_2_), carbon dioxide production (VCO_2_), respiratory exchange ratio (RER), energy expenditure, and total activity as previously described^44^. Mice were placed into individual cages with sensors that measure the food intake. VCO_2_ and VO_2_ levels and energy expenditure were analyzed for four days. Locomotor activity was recorded via breaks of light beams.

### Lipoprotein profile and lipid measurements

To analyze the plasma samples, blood was collected from overnight-fasted mice via the tail vein, and plasma was separated by centrifugation at 10000 rpm at 40°C for 10 minutes. Plasma TAGs were enzymatically analyzed with commercially available kits (Wako Pure Chemicals). The distribution of lipoproteins in the plasma lipid fractions was analyzed by FPLC gel filtration, using 2 Superose 6 HR 10/30 columns (Pharmacia Biotech).

### Hepatic VLDL-TAG secretion

To measure the liver VLDL-TAG secretion rate, mice were fasted overnight and then administered an intraperitoneal injection of 1 g/kg of body weight poloxamer 407 (Sigma-Aldrich) dissolved in PBS. Blood samples were collected from the tail immediately before the injection and at 1, 2, 3, and 4 hours post-injection, following previously described methods^44^. The circulating TAG levels were analyzed using commercially available kits.

### Liver histology and lipid measurement

The liver samples were analyzed by preparing them for histological examination and lipid content determination. For histological analysis, the liver tissues were fixed in 4% paraformaldehyde (PFA), sectioned, processed into paraffin blocks, and stained with H&E. To visualize neutral lipids, frozen liver samples were embedded in OCT, sectioned, and stained with oil red O using the Oil Red O staining method. The total TAG levels were extracted using a chloroform/methanol solvent (2:1) based on a previously described method (reference), and the liver TAG content was measured using a commercially available assay kit (Sekisui) according to the manufacturer’s instructions.

### ALT and AST and AFP measurements

ALT and AST activity and plasma AFP levels were determined in plasma with the commercially assay kits (Sigma-Aldrich, MAK052; and MAK055) following manufacturer’s recommendations.

### Circulating leukocyte analysis

Blood was collected via tail vein in heparinized microhematocrit capillary tubes, and the total numbers of circulating blood leukocytes were analyzed using the HEMAVET system. For further characterization of leukocytes, FACs analysis was performed as follows. Erythrocytes were lysed using ACK lysis buffer (155 mM ammonium chloride, 10 mM potassium bicarbonate, and 0.01 mM EDTA, pH 7.4), and leukocytes were blocked with 2 μg/ml of FcgRII/III. The leukocytes were then stained with a mixture of antibodies, and monocytes were identified as CD115hi and subsets as Ly6-Chi and Ly6-Clo. Neutrophils were identified as CD11bhiLy6Ghi, B cells were identified as CD19hiB220hi, and T cells were identified as CD4hi or CD8hi. The following antibodies were used for all the analysis of leukocytes (all from BioLegend): FITC-Ly6-C (HK1.4), PE-CD115 (AFS98), APC-Ly6-G (1A8), PB-CD11b (M1/70), APC-CD19 (6D5), PE/Cy7-B220 (RA3-6B2), APC/Cy7-CD4 (RM4-5), and BV421-CD8a (53-6.7). All antibodies were used at 1:300 dilutions.

### Isolation of primary hepatocytes and non-parenchymatous cells and hepatic Immune cell Flow Cytometry analysis

Hepatic cells were isolated from the liver of both WT and N-LKO mice obtained from the Yale Liver Facility Center, as previously described (Multistep mechanism of polarized Ca2+ wave patterns in hepatocytes^45^. Briefly, mice were anesthetized with Isoflurane and attached to a Styrofoam tray. The abdomen was wet with 70% ethanol and opened along the midline, and a ligature was placed around the mid-portal vein and IVC. The portal vein was cannulated with a 22 G catheter and perfused with Hanks A followed by Hanks B with collagenase. After the liver was removed, hepatocytes were released by shaking and the cell suspension was filtered through a 40 μm mesh. The cells were then pelleted by centrifugation, and the supernatant was aspirated. The cell pellet was resuspended in L-15 media for further use. The suspension was subjected to centrifugation at 60 g for two minutes to separate pellets containing hepatocytes and non-parenchymatous cells (NPCs). NPCs, which include immune cells, were isolated by centrifuging the suspension at 300 g for 5 minutes and resuspended in 200 μl of ACK solution (155 mM ammonium chloride, 10 mM potassium bicarbonate, and 0.01 mM EDTA, pH 7.4). The NPCs were then stained with a mixture of antibodies to identify specific cell types. B cells were identified using APC-Cy7 B220 (Biolegend, USA), while T cells were identified using CD4hi or CD8hi with the following antibodies: BUV395 CD90.2 - 565257 (BD, USA), BV711 CD4 - 100447 (Biolegend), BV605 CD8a - 100744 (Biolegend). T cell activation was determined by CD62L/CD44 status using PE-Cy7 CD62L - 25-0621-82 (eBioscience, USA), BUV737 CD44 - 612799 (BD), BV605 CD8a - 100744 (Biolegend). Macrophages were identified using FITC F480 - 157310 (Biolegend), and neutrophils were identified using Pacific Blue CD11b - 101224 (Biolegend) and APC Ly6G - 127614 (Biolegend). All antibodies were used at a 1:300 dilution.

### Cis-Prenyltransferase assay (Cis-PTase)

To assay cis-PTase activity in mammalian cells, a reaction mixture was prepared consisting of 250 μg microsomal protein, 25 μM FPP, 50 μM [1-14C]-isopentenyl pyrophosphate (IPP) (55 mCi/mmol), 25 mM Tris-HCl (pH 7.4), 1 mM MgCl2, 1.25 mM DTT, 2.5 mM sodium orthovanadate, 10 μM Zaragozic acid A, and 0.35% Triton X-100 in a total volume of 0.1 ml. The reaction was performed at 37°C for 2 hours and terminated by the addition of 10 μL of concentrated hydrochloric acid. The polyprenol diphosphates were chemically dephosphorylated by incubating the lipids at 90°C for 1 hour. The products were extracted with 4 ml chloroform:methanol (3:2) and washed three times with 1/5 volume of 10 mM EDTA in 0.9% NaCl. The chloroform was evaporated under a stream of nitrogen, and the dephosphorylated lipids were loaded onto HPTLC RP-18 precoated plates and run in acetone containing 50 mM H3P04. The plates were then exposed to film to visualize the products of IPP incorporation. To measure the incorporation of radioactive IPP into the polyprenol fraction, the gel from the zone containing radiolabeled polyprenols was scraped and subjected to liquid scintillation counting. All reagents used were of analytical grade and purchased from Sigma-Aldrich, Thermo Fisher Scientific, and [1-14C] IPP (50 mCi/mmol) was purchased from American Radiolabeled Chemicals. The reverse-phase TLC (RP18-HTLC) plates used were from MilliporeSigma (catalog no. 1.16225.0001).

### Western blot analysis

To prepare the liver homogenates, we utilized the Bullet Blender Homogenizer method as previously described. Tissues were lysed in an ice-cold buffer containing 50 mM Tris–HCl (pH 7.5), 0.1% SDS, 0.1 mM EDTA, 0.1% deoxycholic acid, 0.1 mM EGTA, 1% NP-40, 5.3 mM NaF, 1.5 mM Na4P2O7, 1 mM orthovanadate, 1 mg/ml protease inhibitor cocktail (Roche), and 0.25 mg/ml AEBSF (Roche). The lysates were sonicated and rotated at 4°C for one hour, followed by centrifugation at 12,000 g for 30 minutes.

After normalizing the protein concentration, equal amounts of proteins were resuspended in SDS sample buffer and separated by SDS-PAGE. The separated proteins were then transferred onto nitrocellulose membranes and probed with various antibodies such as anti-ATF4, Bip1and anti-HSP90. The protein bands were detected using the Odyssey Infrared Imaging System (LI-COR Biotechnology), and densitometry was performed using ImageJ software.

For the immunoblot analysis of ApoB-100 and ApoB-48 in pooled VLDL lipoprotein fractions, separation was performed using a NuPAGE Novex 3%–15% Tris-Acetate Mini Gel with 1× NuPAGE Tris-Acetate SDS running buffer (Invitrogen). Following an overnight transfer of proteins onto nitrocellulose membranes, the membranes were blocked with nonfat milk dissolved in wash buffer 5% (w/v). The membrane was then probed with an antibody against ApoB (Meridian, K23300R; 1:2,000) overnight at 4°C.

### Comparative Gene Expression Profiling in the Liver of NASH Patients and Healthy Humans

To investigate the expression of Nus1 in human liver with nonalcoholic steatohepatitis (NASH) compared to healthy liver, we obtained RNA-seq gene expression data from a publicly available human liver cohort (GSE135251) through the Gene Expression Omnibus (GEO). Processed data files were downloaded from GEO (data processing is outlined in (https://www.science.org/doi/10.1126/scitranslmed.aba4448)), then normalized and transformed using edgeR-limma-voom as described (https://www.ncbi.nlm.nih.gov/pmc/articles/PMC4937821/, https://genomebiology.biomedcentral.com/articles/10.1186/gb-2014-15-2-r29). The results were presented as box-and-whisker plots, with the central lines representing medians, the edges of the box indicating upper and lower quartiles, and the whiskers indicating the minimum and maximum values.

### RNA isolation and quantitative real-time PCR

The isolation of total RNA from tissue or cells was conducted using TRIzol reagent (Invitrogen) in accordance with the manufacturer’s instructions. Subsequently, cDNA was synthesized using iScript RT Supermix (Bio-Rad) as per the manufacturer’s protocol for mRNA expression analysis. For quantitative real-time PCR (qRT-PCR) analysis, Sso Fast Eva Green Supermix (Bio-Rad) was utilized, and the measurements were performed on an iCycler Real-Time Detection System (Eppendorf). The mRNA levels were normalized to 18S.

### Measurement of ROS generation

The cellular ROS species H2O2 and O2- were analyzed in primary hepatocytes from WT and N-LKO mice using DCFDA and DHE dyes, following the manufacturer’s instructions. Firstly, isolated hepatocytes were incubated with 5 μM of DCFDA or DHE dyes for 30 minutes at 37°C. Next, the cells were washed twice with PBS and their fluorescence was acquired using flow cytometry (FACS Area, BD Bioscience).

### Deep RNA seq

Total RNA was isolated and purified from both the livers of control and N-LKO mice or tumors of N-LKO mice using a RNA isolation Kit from Qiagen, followed by DNAse treatment to eliminate any genomic contamination using RNA Min Elute Cleanup from Qiagen. The purity of the total RNA samples was confirmed using the Agilent Bioanalyzer from Agilent Technologies.

The RNA sequencing was performed by the Yale Center for Genome Analysis (YCGA) after the removal of rRNA from RNA samples using the Ribo-Zero rRNA Removal Kit from Illumina. The RNA libraries were created using the TrueSeq Small RNA Library preparation kit from Illumina, and then sequenced for 45 cycles on the Illumina HiSeq 2000 platform (1 x 75bp read length). To ensure high quality of the reads, in-house developed scripts were used to trim for quality, and the reads were aligned to the reference genome using TopHat2. Transcript abundances and differences were calculated using cuffdiff, and the results were analyzed using R and cummeRbund through in-house developed scripts. The RNA sequencing data has been deposited in the Gene Expression Omnibus database (GSE230972).

### Whole genome sequencing and data analysis

We extracted genomic DNA (gDNA) samples from the liver and tumor tissues of N-LKO mice fed with a Western diet (WD) using the Promega Reliaprep gDNA Tissue Miniprep System following the manufacturer’s protocol. We used 0.5 µg DNA for library preparation, which involved enzymatic fragmentation, end-repair, and A-tailing in a single reaction using the xGen™ DNA EZ Library Prep Kit (IDT, Part#10009021). We then ligated the appropriate dual multiplexing indices, xGen UDI-UMI Adapters (IDT, Part #10005903), to the DNA fragments for hybridization to the flow-cell for cluster generation. We determined the size of the final library construct on the Caliper LabChip GX system and quantified it using qPCR SYBR Green reactions with a set of DNA standards and the Kapa Library Quantification Kit (KAPA Biosystems, Part#KK4854). Size and concentration values were entered into the WikiLIMS database for the sequencing team’s use for appropriate flow-cell loading. For flow-cell preparation and sequencing, we normalized sample concentrations to 2nM and loaded them onto Illumina NovaSeq 6000 flow cells at a concentration that yields at least 700Gbp of passing filter data per lane. We optimized the loading concentration for whole-genome sequencing (WGS) libraries to maximize both well occupancy and unique read output while limiting duplicates associated with patterned flow-cell technology. We sequenced samples using 151 bp paired-end sequencing reads according to Illumina protocols. The 10bp indexes were read during additional sequencing reads that automatically followed the completion of read 1. Data generated during sequencing runs were simultaneously transferred to the YCGA high-performance computing cluster. A positive control, a prepared bacteriophage Phi X library provided by Illumina, was spiked into every lane at a concentration of 1% to monitor sequencing quality in real-time. For data analysis, signal intensities were converted to individual base calls during a run using the system’s Real Time Analysis (RTA) software. Base calls were then transferred from the machine’s dedicated personal computer to the Yale High-Performance Computing cluster via a 1 Gigabit network mount for downstream analysis. Primary analysis, including sample de-multiplexing and alignment to the human genome, was performed using Illumina’s CASAVA 1.8.2 software suite.

To perform variant calling on the sequencing reads, we followed the GATK 4 best practices guidelines^46^. The reads were aligned to the mm10 mouse reference using BWA MEM^47^, and duplicates were marked using picard’s Mark Duplicates command. Base quality scores were recalibrated, and aligned CRAM files were generated using GATK. Joint variant calling of the two samples was performed using Haplotype Caller and Genotype GVCFs. Variants were hard-filtered using GATK’s Select Variants and Variant Filtration with their best practice settings. VCF files were annotated with Variant Effect Predictor (VEP)^48^. A final variant list was created by filtering passed variants whose genotype differed between the two samples. Mutational counts were computed from this variant list, and the mutational signatures were visualized using the sig Profiler Plotting package^49^.

### Single cell RNA sequencing and data analysis

We performed single-cell RNA sequencing (scRNA-seq) on livers using the 10x Genomics Chromium platform. Hepatic cells were isolated from both wild-type (WT) and N-LKO mice obtained from the Yale Liver Facility Center. We used a multi-step mechanism for isolating hepatocytes from the liver, which has been described previously^45^. Briefly, the mice were anesthetized with Isoflurane and attached to a Styrofoam tray. The abdomen was wet with 70% ethanol, and a ligature was placed around the mid-portal vein and IVC. The portal vein was cannulated with a 22 G catheter and perfused with Hanks A followed by Hanks B with collagenase. The liver was then removed, and the hepatocytes were released by shaking. The cell suspension was filtered through a 40 μm mesh, and the cells were pelleted by centrifugation. The cell pellet was resuspended in L-15 media for further use.

We mixed hepatocytes and non-parenchymatous cells (NPCs) in a 1:1 ratio for WT mice and hepatocytes, NPCs, and tumor cells in a 1:1:0.25 ratio for N-LKO mice. We loaded single cells onto the 10x Genomics Chromium Single-Cell controller at the Yale Center for Genome Analysis, followed by lysis and barcoded reverse transcription of polyadenylated mRNA from each cell using the Single Cell 3’ Reagent Kits v3. The libraries were sequenced on an Illumina HiSeq 4000 as 2 × 150 paired-end reads.

We used Cell Ranger software for sample demultiplexing, read alignment, and unique molecular identifier (UMI) processing. Low-quality cells, doublets, and potentially dead cells were filtered out based on the percentage of mitochondrial genes and number of genes and UMIs expressed in each cell.

To cluster the cells, we used the Seurat R package with filtered genes by barcode expression matrices as inputs. We calculated highly variable genes (HVGs) using the Seurat function "Find Variable Genes" and used them for downstream clustering analysis. We visualized clustering of liver single-cell transcriptomes (7000 cells from WT and 7000 cells from N-LKO mice fed WD) using uniform manifold approximation and projection (UMAP). UMAP was performed with the "Run UMAP" function (Seurat) using HVGs for dimensionality reduction. Clustering was done through the "Find Clusters" function using 30 significant PCs with a resolution of 0.3. We also identified significantly differentially expressed genes in a cluster using the Seurat function "Find All Markers" expressed in more than 25% of cells with at least 0.25-fold difference and reaching statistical significance of an adjusted p<0.05 as determined by the Wilcox test.

Finally, the scRNA-seq data generated in this study were deposited in NCBI Gene Expression Omnibus (GSE230972).

### Statistics

The mouse sample size for each study was based on literature documentation of similar well-characterized experiments. The number of mice used in each study is listed in the figure legends. No inclusion or exclusion criteria were used, and studies were not blinded to investigators or formally randomized. Data are expressed as average ± SEM. Statistical differences were measured using an unpaired 2-sided Student’s t-test, or 1-way or 2-way ANOVA with Bonferroni’s correction for multiple comparisons. Normality was tested using the Kolmogorov-Smirnov test. A nonparametric test (Mann-Whitney) was used when the data did not pass the normality test. A value of P ≤ 0.05 was considered statistically significant. Data analysis was performed using GraphPad Prism software version 7.

## Acknowledgments

This research was supported by the National Institutes of Health (N.I.H.) through grants R35HL139945, RO1DK125492, and American Heart Association MERIT Award (W.C.S.), an American Heart Association Postdoctoral Fellowship Award (S.L.), and N.I.H. grant K01DK124441 (N.E.B.).

## Authors contributions

A.K.S. and W.C.S. conceived and designed the study, as well as wrote the manuscript. A.K.S. conducted the majority of the experiments and analyzed the data. B.C. performed experiments, analyzed the single-cell RNA sequencing data, and contributed to manuscript editing. K.M.C. and J.W.F. analyzed the deep RNA sequencing data. K.H. isolated the liver cells from the mice. M.S. and J.C. conducted the metabolic cage experiment. I.R.M. ran the FPLC for lipoprotein profiling. K.G. measured the cis-PTase enzyme activity. S.W. and S.L. conducted experiments and analyzed the data. S.S. and N.B. conducted experiments, analyzed data, and contributed to manuscript editing. TTR provided the DGAT2 inhibitor. J.K. analyzed whole-genome sequencing data.

---

**Extended Data Fig. 1.**
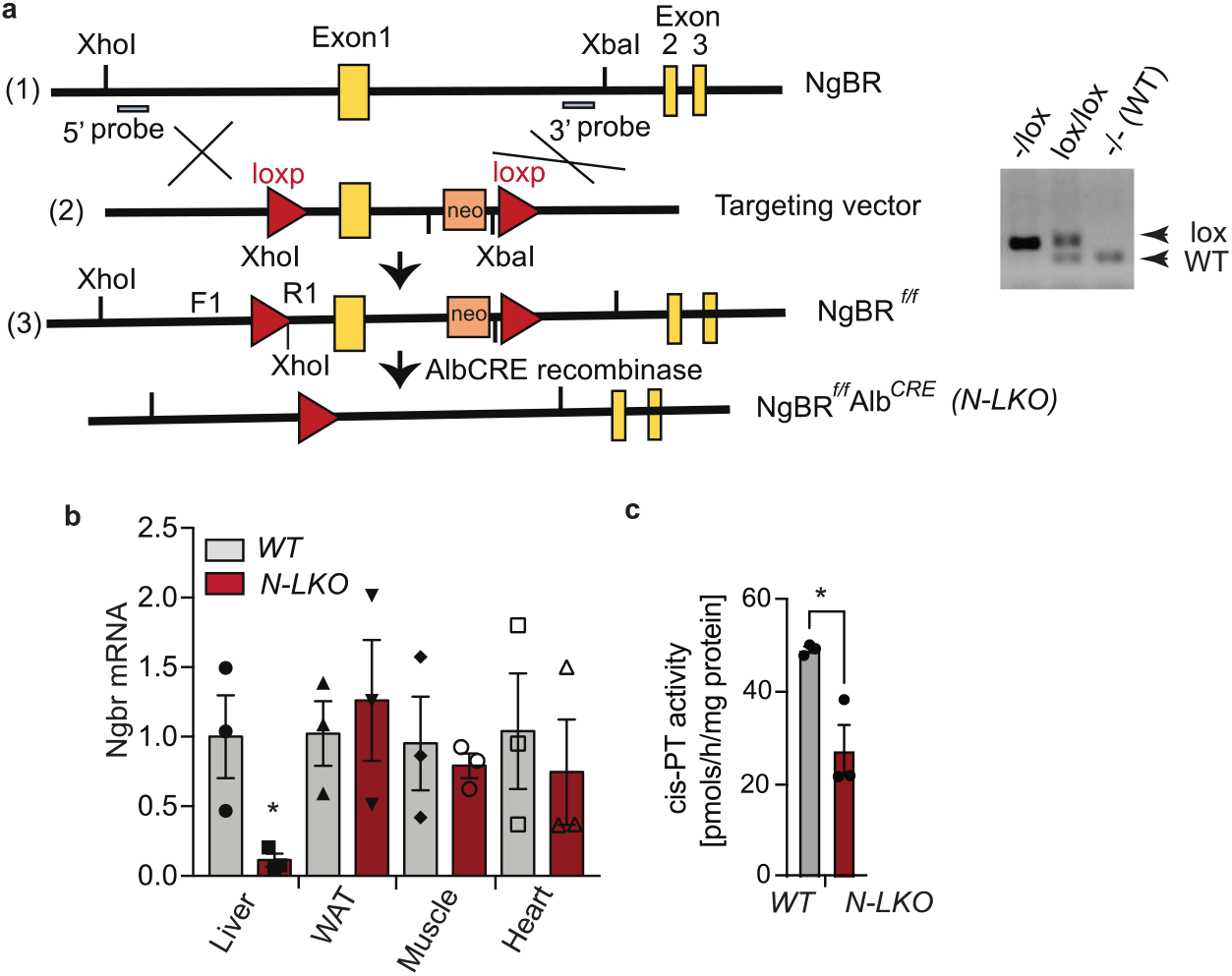
The Loss of NgBR function in the liver reduces cis-PTase enzyme activity. **a,** Schematic diagram showing the generation of liver-specific NgBR knockout (N-LKO) mice. (1) The NgBR genomic DNA fragment is composed of three exon 1-3 (2) Schematic construct of the NgBR targeting vector. NgBR exon1 is flanked by two loxP sites. (3) Mice with floxed allele are generated after homologous recombination. Consequently, these mice were bred with mice expressing CRE recombinase to generate tissue-specific NgBR-knockout mice. Right panel represents PCR amplification of NgBR*^f/f^* mice showing bands from both, one, or none of the floxed alleles. mRNA expression of NgBR in the liver, WAT, heart and muscle of WT and N-LKO mice (n=3). **b,** NgBR expression in these tissues is normalized to its expression in liver from WT mice. **c,** Measurement of microsomal Cis-PTase activity in isolated hepatocytes of WT and N-LKO mice (n=3). Two-sided \**P* < 0.05; comparing N-LKO with WT mice using an unpaired Welch’s t-test.

**Extended Data Fig. 2.**
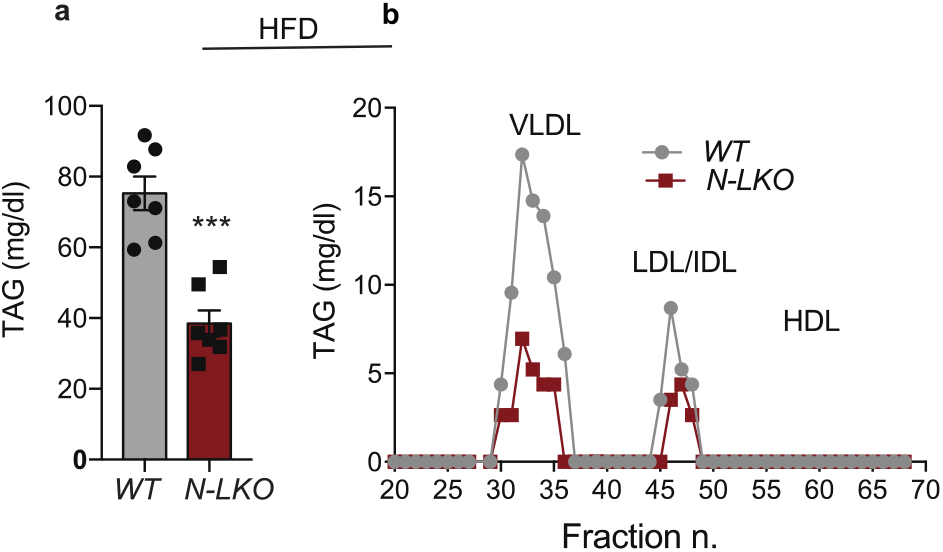
The hepatic loss of NgBR function reduces circulating TAG levels under obesogenic diet conditions. **a,** Circulating TAG levels from overnight fasted WT and N-LKO mice fed high-fat-diet (HFD) for 4 months (n=7). **b,** TAG content of FPLC-fractionated lipoproteins from pooled plasma (n=5) of overnight-fasted WT and N-LKO mice fed a HFD for 4 months. Two-sided ***P < 0.001, by Welch’s t-test.

**Extended Data Fig. 3.**
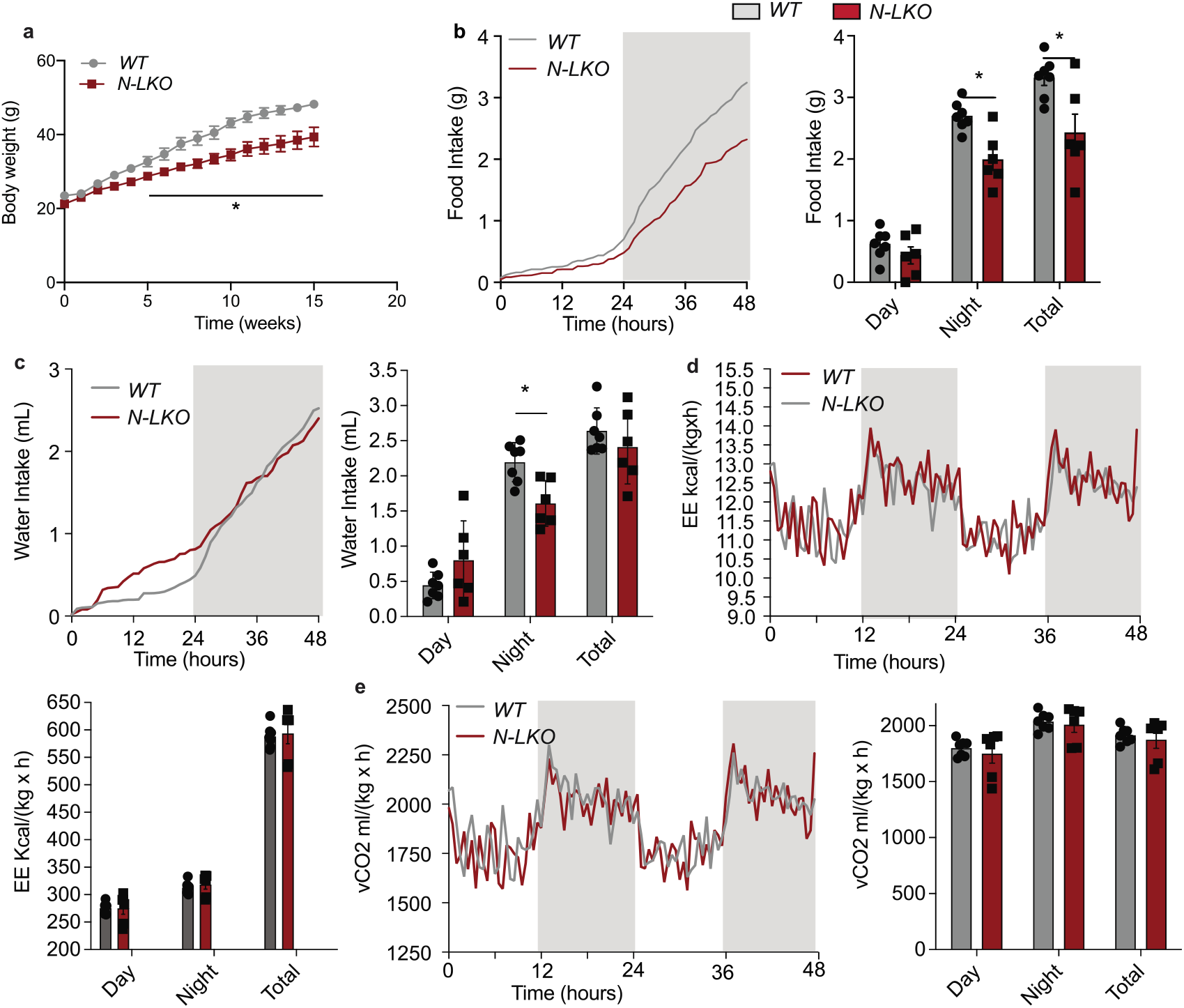
Suppression of hepatic NgBR reduces body weight food consumption and water intake without influencing energy expenditure. **a,** Body weight of WT and N-LKO mice fed HFD for 16 weeks. **b-e,** Metabolic analysis of HFD fed WT and N-LKO mice over 48hour period. **b,** Food consumption. Cumulative graph on the right**. c,** Water intake. Right panel displays cumulative graph of water intake. **d,** Energy expenditure (EE) Right section shows cumulative graph. **e,** Carbon dioxide production. Cumulative graph on the right panel. All data are represented as mean ± SEM. \**P* < 0.05; comparing N-LKO with WT mice using an unpaired Welch’s *t* test, (n=7).

**Extended Data Fig. 4.**
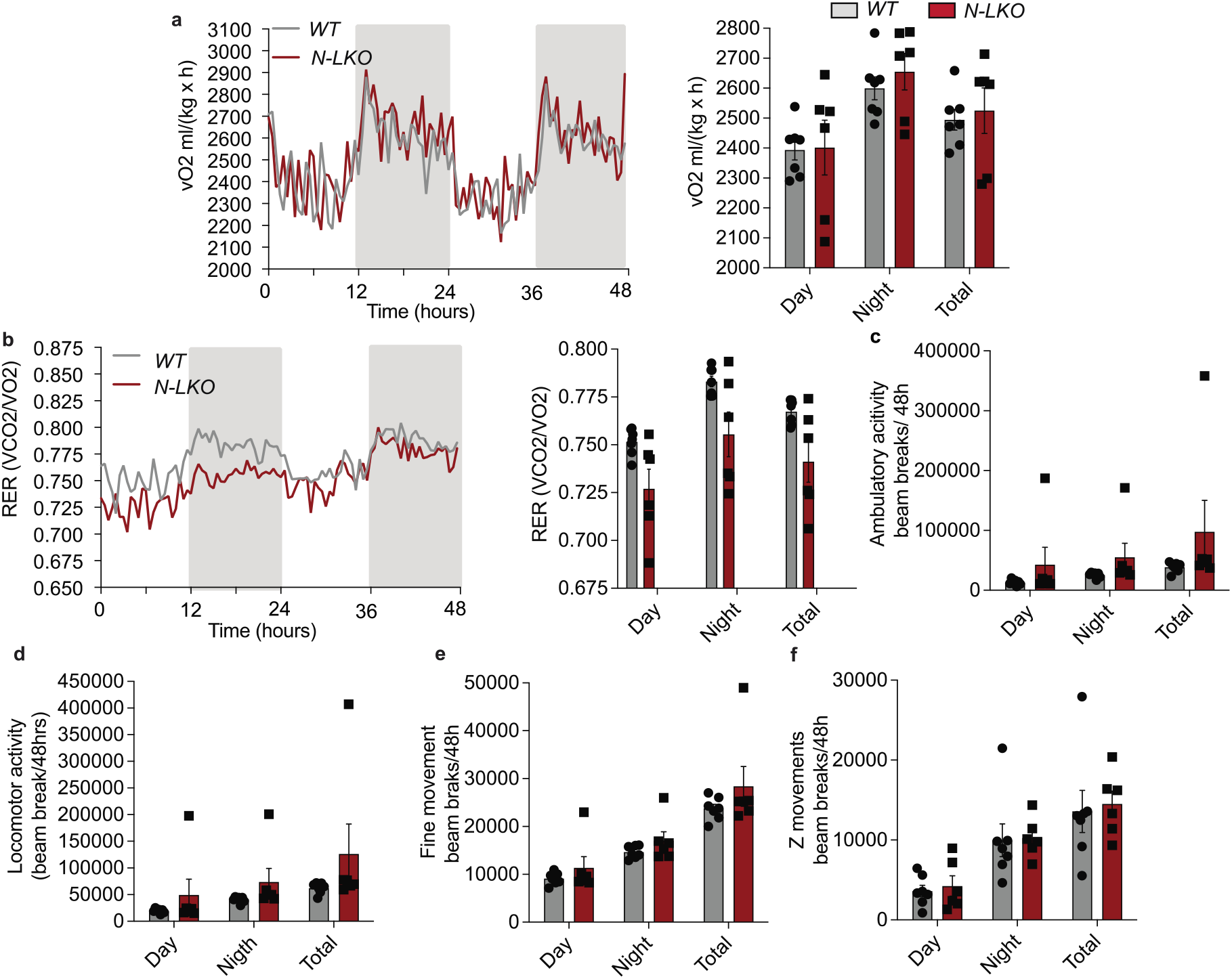
Ablation of hepatic NgBR does not affect the locomotor activity. Metabolic analysis of HFD fed WT and N-LKO mice over 48hour period. **a,** Oxygen consumption. Right panel shows cumulative graph. (n=7). **b,** Respiratory exchange ratio (RER) **c,** Ambulatory activity, **d,** Locomotor activity, **e,** fine movement and **f,** Z Movement (n=7).

**Extended Data Fig. 5.**
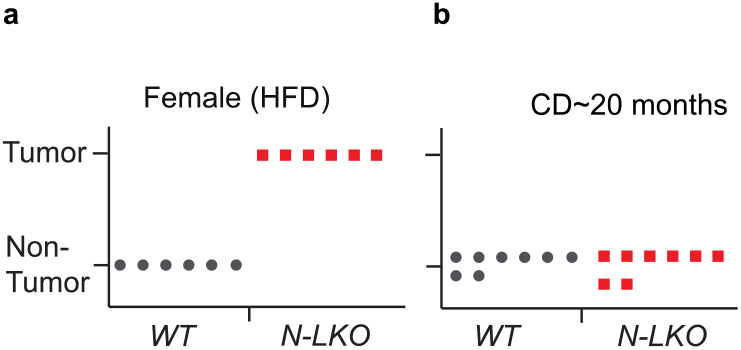
Hepatic loss of NgBR drives the HCC development in HFD- diet conditions. **a,** The graph show tumor incidence in female N-LKO mice fed an HFD for 16 weeks. **b,** The graph show no tumor incidence in N-LKO mice fed an CD for ∼ 20 months.

**Extended Data Fig. 6.**
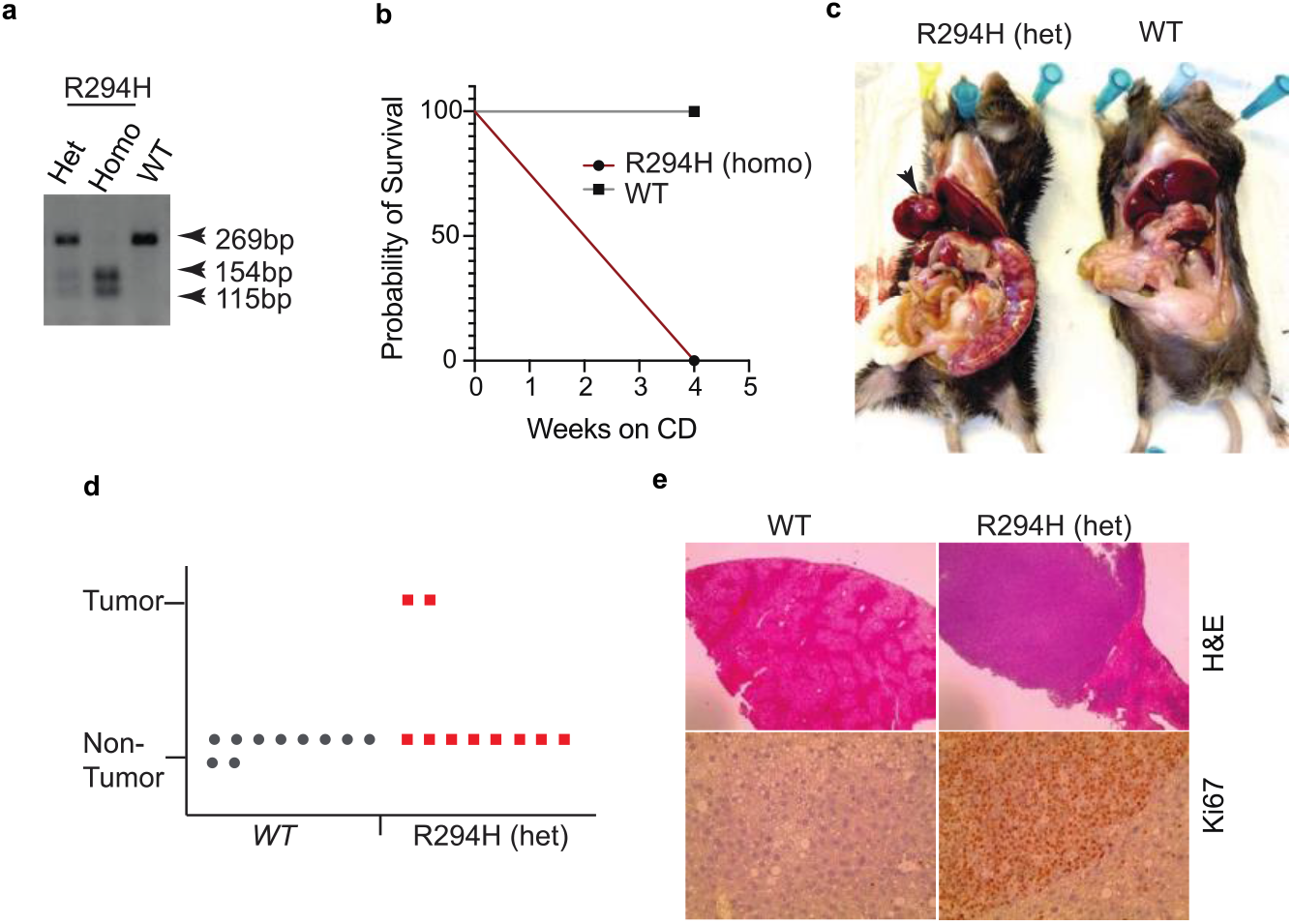
Mutation in NgBR drives the HCC development in HFD- diet conditions. **a,** Generation of NgBR R-294H mutant mice. PCR and agarose gel electrophoresis of WT, R294H point mutant homozygous and heterozygous tail samples. Genotyping from *NgBR R294H mutant* mice presentation bands from one (WT,269bp) two (homozygous-154+115bp) or three (heterozygous -269 and 154+115bp). **b,** Kaplan-Meier survival curves of WT and R294H mutant homozygous mice. **c,** Representative image of mice with liver cancer of R294H heterozygous mutant and WT fed HFD for 16 weeks. arrow showing toward HCC. **d,** Right panel shows graph summarizing R294H mutant heterozygous and WT mice with or without tumor on HFD (n=8). Symbols display individual mice. **e,** Histological analysis of liver and tumor sections stained with H&E and Ki-67 isolated from WT and R294H mutant heterozygous mice on HFD. Scale bar, 200 μm.

**Extended Data Fig. 7.**
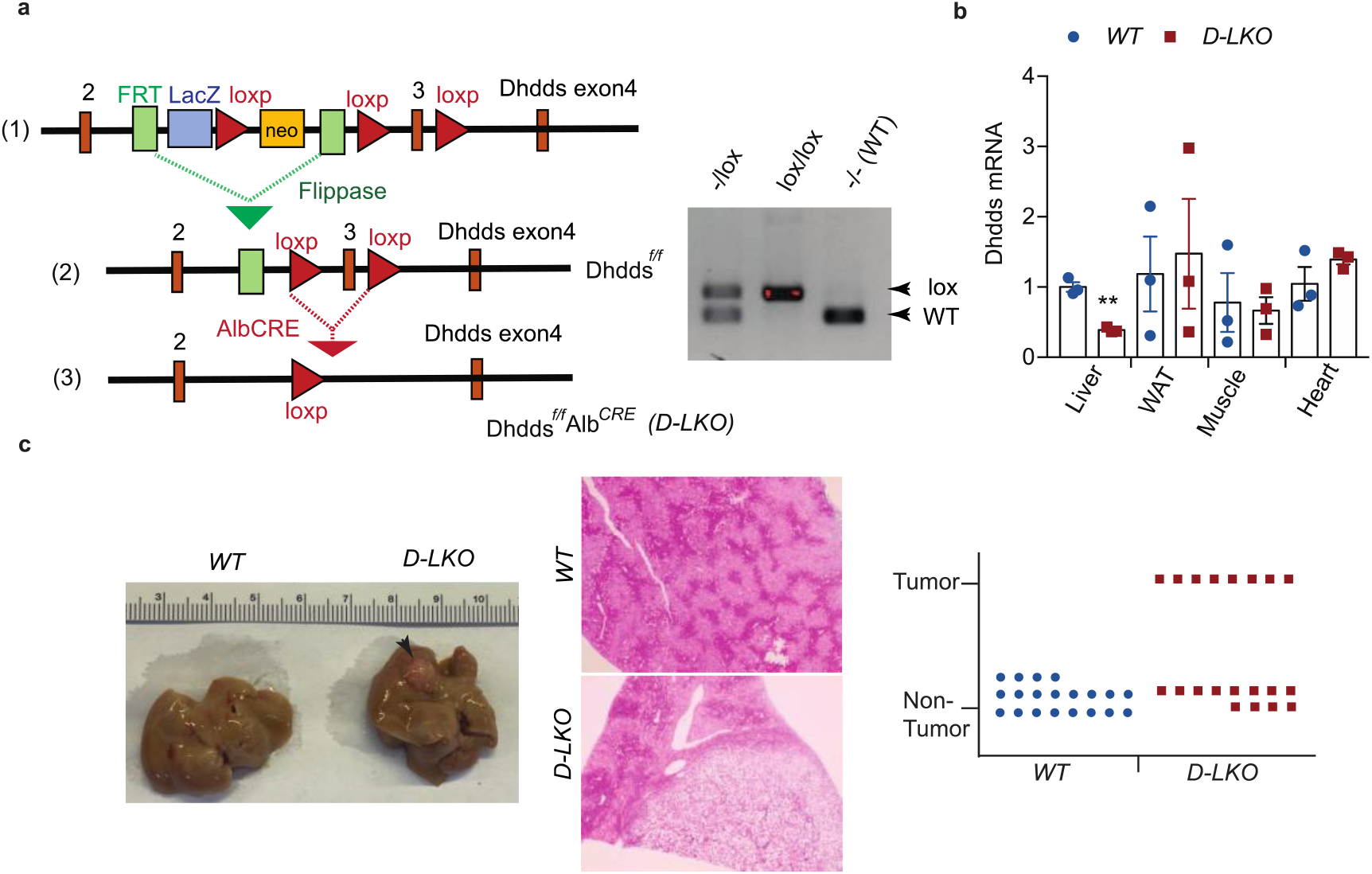
Depletion of Dhdds in the liver drives the HCC development in HFD-diet conditions. **a,** Schematic diagram showing the generation of liver-specific Dhdds knockout (D-LKO) mice. (1) The cassette is composed of a short flippase recombination enzyme (Flp) recognition target (FRT), a Cre recombinase recognition target (loxP). Dhdds exons 2-3 are flanked by the loxP site. (2) Mice with floxed allele but missing neomycine cassette were generated by crossing with flp recombinase-deleter mice. (3) Afterward, these floxed mice were bred with mice expressing Cre recombinase to generate tissue-specific (D-LKO) mice. Genotyping from *Dhdds^fl/fl^* mice presentation bands from one, both, or none of the floxed alleles. **b,** mRNA expression of Dhdds in the liver, WAT, muscle, and heart of WT and D-LKO mice (n=3). Dhdds expression in the tissue (s) is normalized to its expression in liver from WT mice. **c,** Representative images of the liver isolated from WT and D-LKO mice fed a HFD for 16 weeks, arrow showing toward HCC. Right panel represents histological analysis of liver and tumor sections stained with H&E and right panel shows graph summarizing D-LKO and WT mice with or without tumor feeding on HFD. Symbols represent individual mice. Scale bar, 200 μm. Two-sided; \*\**P* < 0.01; comparing *D-LKO* with WT mice using an unpaired Welch’s t-test.

**Extended Data Fig. 8.**
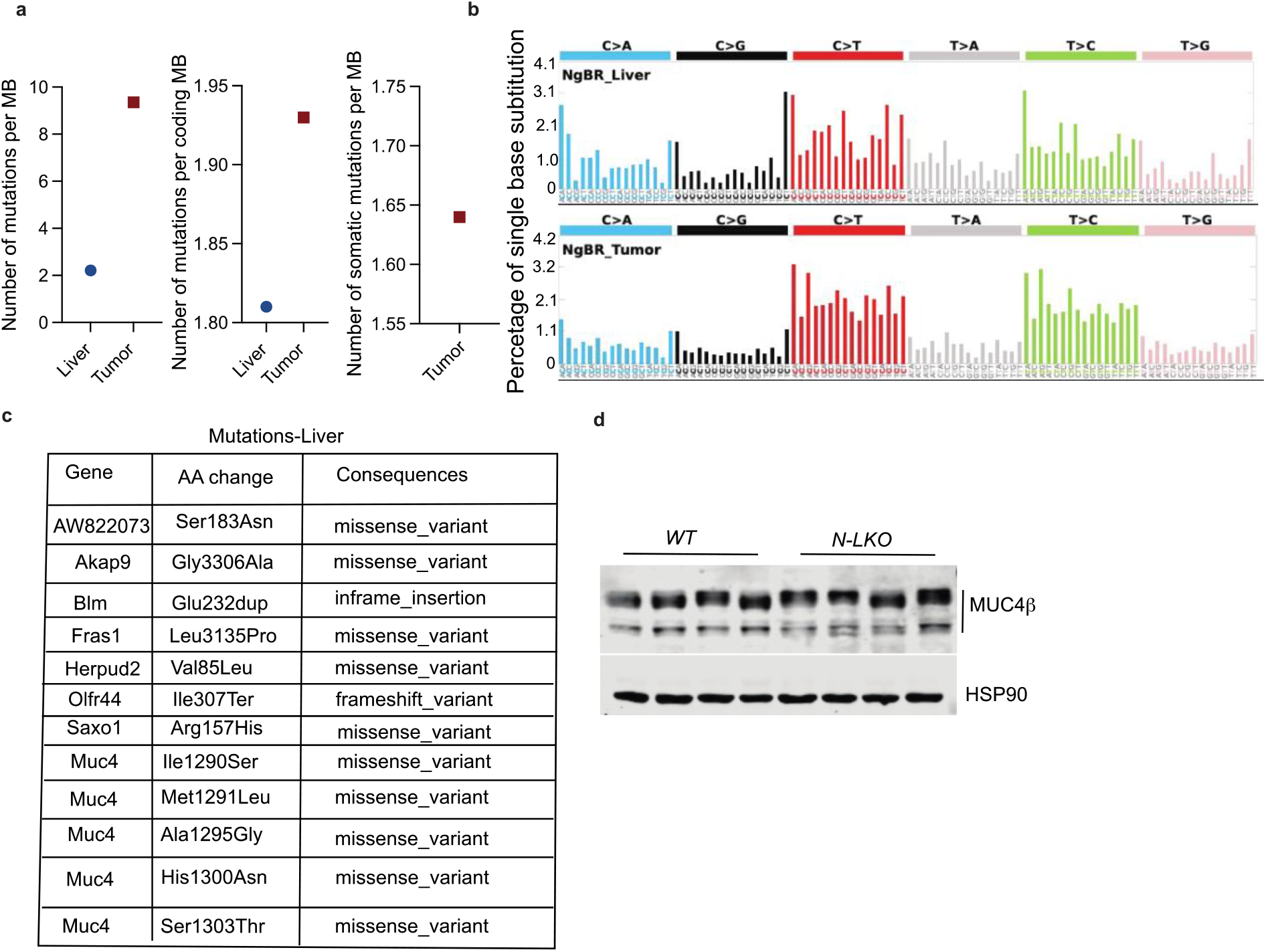
Mutation landscape analysis of liver and tumor initiated by hepatic NgBR deficiency in mouse. **a,** Comparison of mutation burden, measured as the number of mutations per megabase (MB) or per coding MB, in tumors and tumor-adjacent liver tissue, as well as the number of somatic mutations per MB in tumors from N-LKO mice fed a Western diet (WD) for 16 weeks. **b,** The array shows the mutation signature analysis of adjacent liver (upper panel) and tumor (lower panel) from a mouse with hepatic NgBR deficiency fed a western-type diet (WD) for 16 weeks. The x-axis represents the base substitution pattern based on trinucleotide context, while the y-axis shows the percentage of single base substitution. **c,** The table displays the list of gene variants identified in the adjacent liver tissue of N-LKO mice. **d**, Representative western blots of MUC4 and housekeeping standard HSP90 in liver lysate from WT and N-LKO mice fed a WD for 16 weeks (n=4).

**Extended Data Fig. 9.**
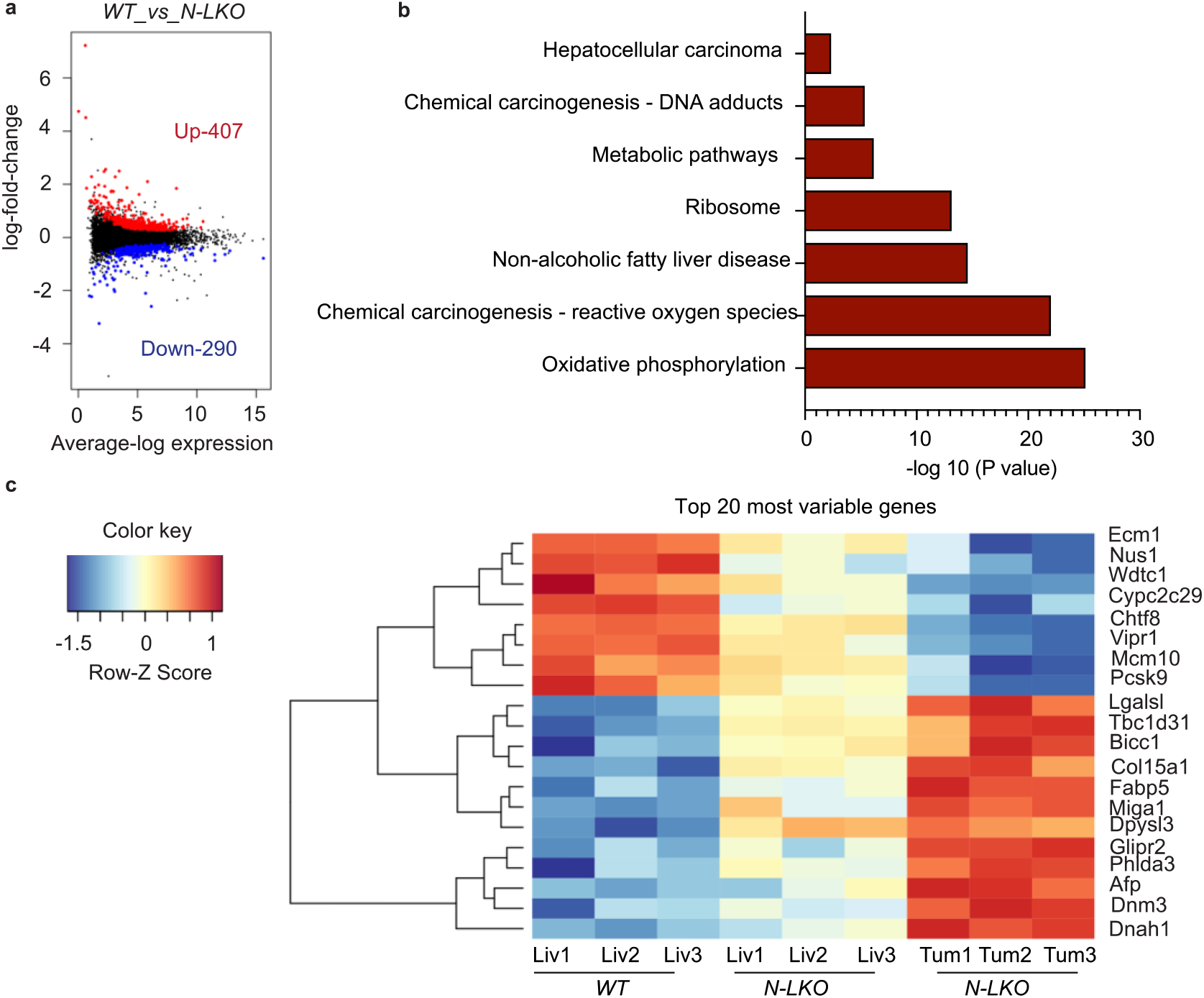
Liver-specific NgBR deficiency enhances the expression of genes related to NAFL and HCC. **a,** RNA seq analysis in the livers of WT and N-LKO mice fed a HFD. MBD plot for differential expression genes that showing the log-fold change and average abundance of each gene. Loss of hepatic NgBR upregulated 407 genes and downregulated 290 genes (n=4). **b,** KEGG pathway analysis of differentially upregulated genes in N-LKO vs WT (n=3). **c,** RNA seq analysis in livers of WT and N-LKO and tumors of N-LKO mice fed a WD (n=3). Heat map representing differentially expressed genes involved in hepatic cancer.

**Extended Data Fig. 10.**
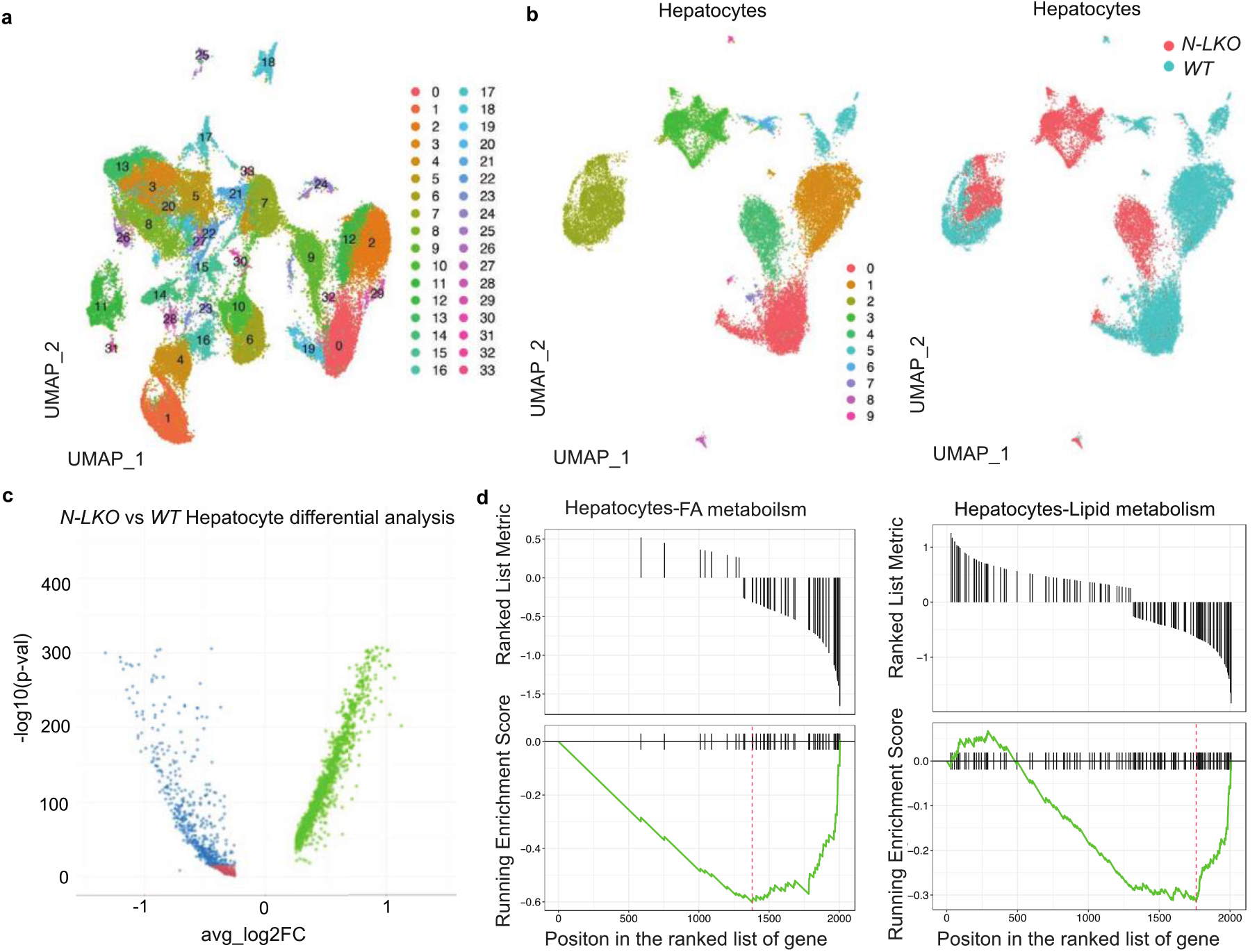
Liver-specific NgBR ablation induces distinct and differential gene expression patterns in liver cells. **a,** Single-cell RNA sequencing analysis of liver cells from WT and N-LKO mice fed a Western diet (WD) UMAP plot showing 33 distinct clusters of cells isolated from the liver of WT and N-LKO mice fed WD. **b,** UMAP plots representing 9 subclusters of hepatocytes in WT and N-LKO mice. The color represents the subcluster or genotype. **c.** Volcano plot displaying the differential expression of genes in log-transformed upregulated and downregulated genes. **d,** Gene set enrichment analysis (GSEA) plot representing the downregulation of genes in hepatocytes involved in fatty acid and lipid metabolism.

**Extended Data Fig. 11.**
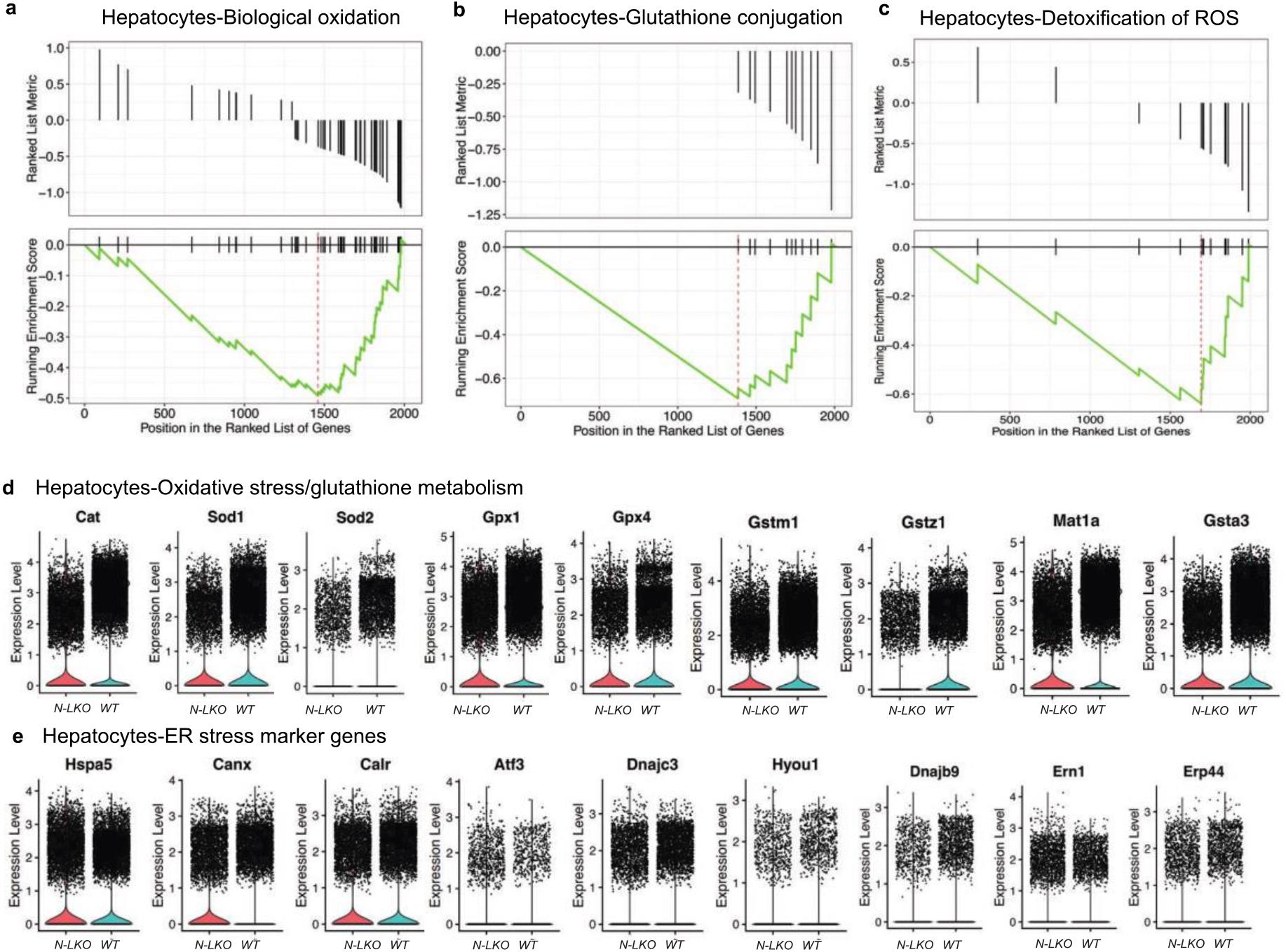
Liver-specific NgBR depletion downregulates oxidative stress response. Single-cell RNA sequencing analysis of hepatic cells from WT and N-LKO mice fed a Western diet (WD**). a-c,** Gene set enrichment analysis (GSEA) plot representing the downregulation of genes involved in biological oxidation, glutathione conjugation and detoxification of ROS (reactive oxygen species) in hepatocytes of N-LKO mice compared to those of WT mice. **d-e,** The violin plots indicate a significant reduction in the expression of genes related to oxidative stress, glutathione metabolism, and ER stress in the hepatocytes of N-LKO compared to WT mice.

**Extended Data Fig. 12.**
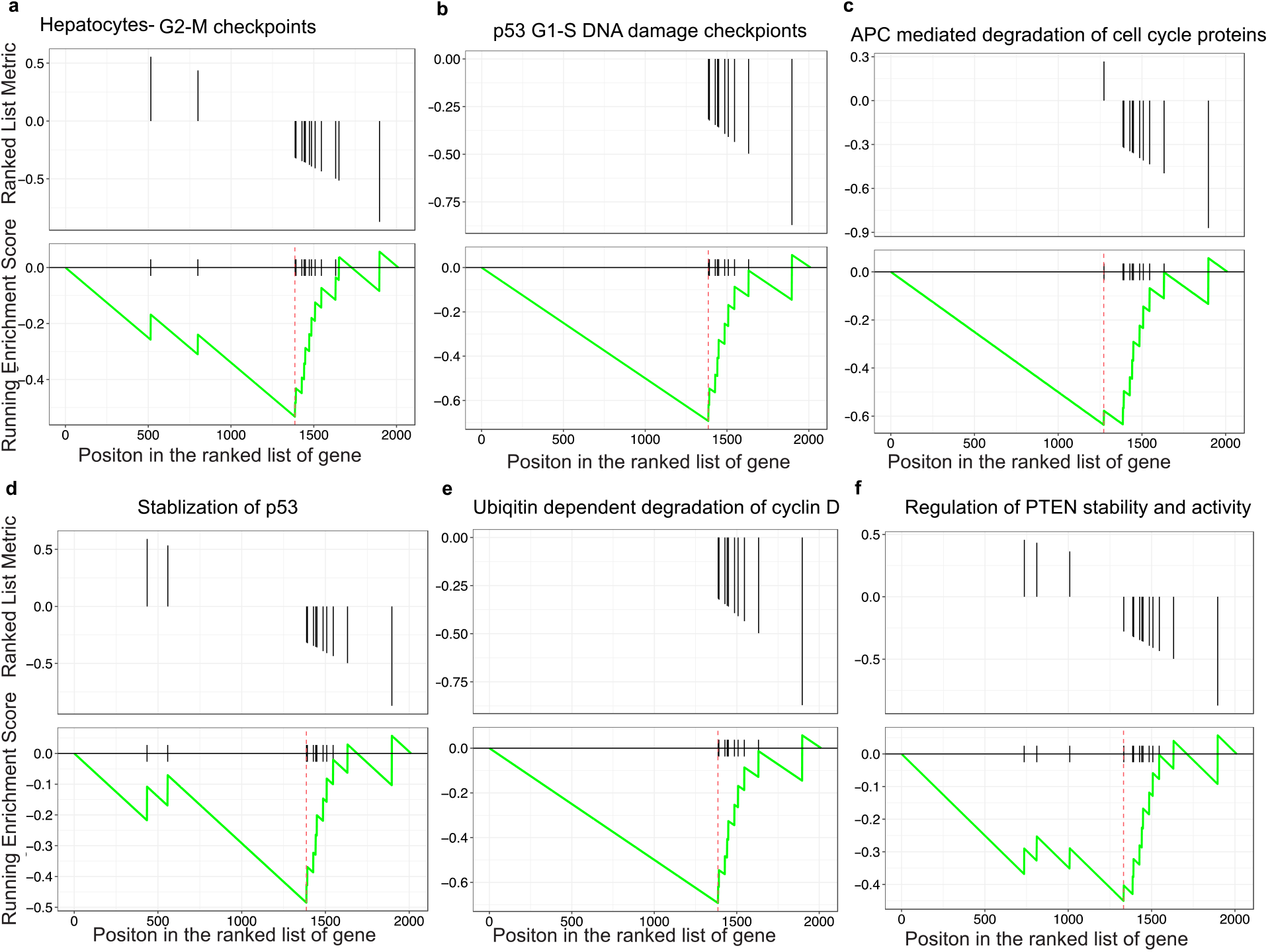
Lack of NgBR reduces the cell cycle checkpoint regulators. Single-cell RNA seq analysis distinct gene expression patterns in the hepatocytes of WT and N-LKO mice fed a Western diet. **a,** Gene set enrichment analysis (GSEA) plot representing genes involved in the G2-M checkpoint regulation are downregulated in the hepatocytes of N-LKO mice compared to WT mice. **b,** Genes involved in p53-dependent G1-S DNA damage checkpoint regulation are significantly downregulated in N-LKO mice compared to WT mice fed a WD. **c,** APC-mediated degradation of cell cycle proteins: genes involved in the APC-mediated degradation of cell cycle proteins are downregulated in the N-LKO mice compared to WT mice. **d,** Stabilization of p53: genes involved in the stabilization of p53, a tumor suppressor protein that regulates the cell cycle, are downregulated in the hepatocyte of N-LKO mice compared to WT mice. **e,** Ubiquitin-dependent degradation of cyclin D: genes involved in the ubiquitin-dependent degradation of cyclin D, a protein that regulates the G1 phase of the cell cycle, are downregulated in the hepatocyte of N-LKO mice compared to WT mice. **f,** Regulation of PTEN stability and activity: genes involved in the regulation of PTEN stability and activity, a tumor suppressor protein that regulates cell cycle progression and cell growth, are downregulated in the hepatocytes of N-LKO mice compared to WT mice. The normalized enrichment score (NES) for each gene set is shown on the plot, with negative NES indicating downregulation of genes in the N-LKO mice compared to WT mice.

**Extended Data Fig. 13.**
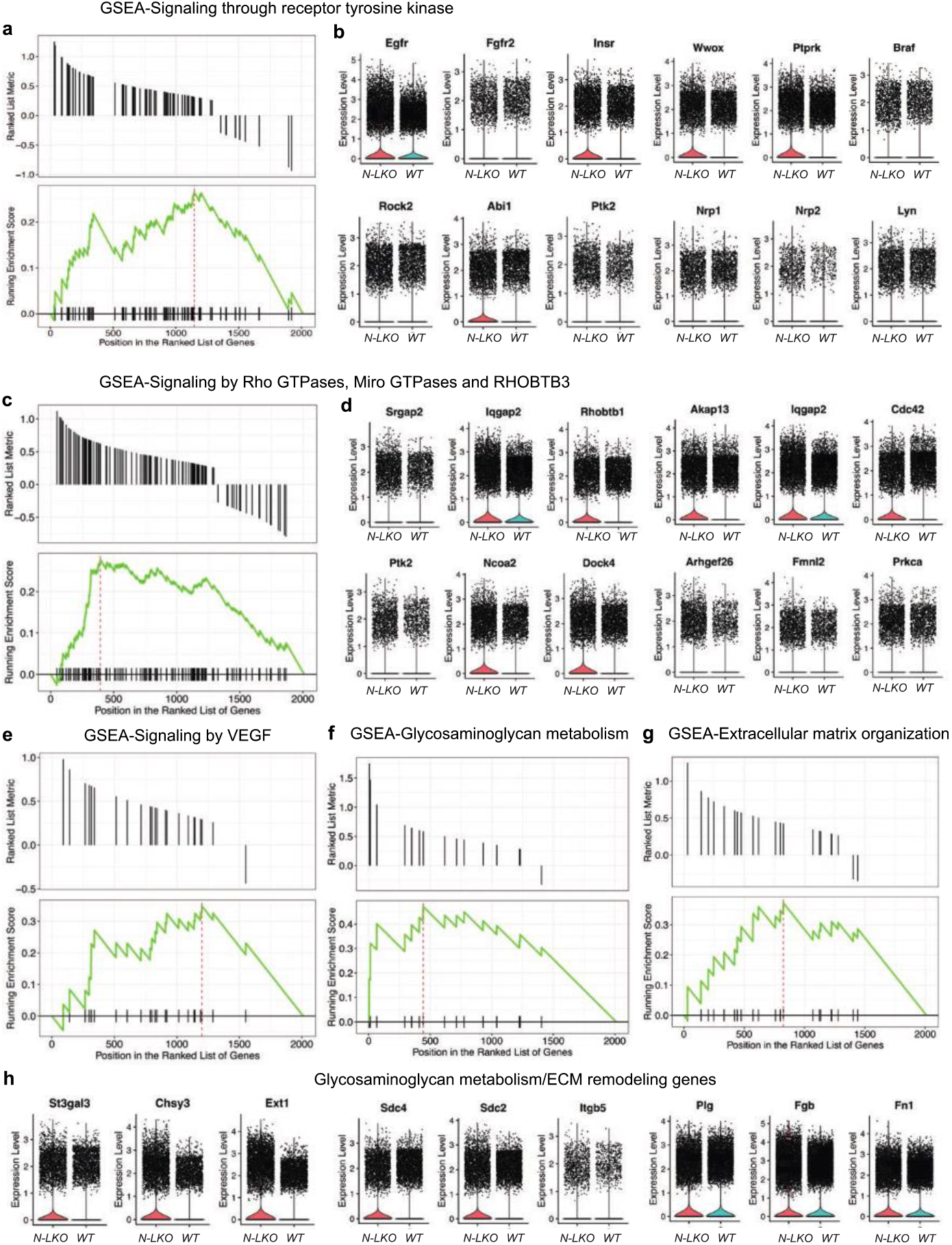
Absence of NgBR in hepatocytes promotes oncogenic associated pathway. Single-cell RNA sequencing analysis that identified distinct gene expression patterns in the hepatocytes of N-LKO mice that were fed a Western diet. **a,** The gene set enrichment analysis (GSEA) plot reveals that genes involved in signaling via tyrosine receptors were upregulated in hepatocytes of N-LKO mice compared to WT mice. **b,** The violin plots further reveal significantly increased expression levels of hepatocyte genes involved in activation of tyrosine receptor signaling in N-LKO mice compared to WT mice. **c,** The GSEA plot shows the upregulation of genes involved in Rac/Rho GTPase signaling in hepatocytes of N-LKO mice compared to WT mice. **d,** The violin plots illustrate expression levels of hepatocyte genes involved in Rac/Rho GTPase signaling were significantly elevated in N-LKO mice relative to WT mice. **e-g** The GSEA plots show the upregulation of genes involved in VEGF signaling, glucosamine glycans metabolism, and ECM organization in hepatocytes of N-LKO mice compared to WT mice. **h,** The violin plots demonstrate a significantly increase in the expression of genes involved in glucosamine glycans metabolism and ECM remodeling in N-LKO mice relative to WT mice.

**Extended Data Fig. 14.**
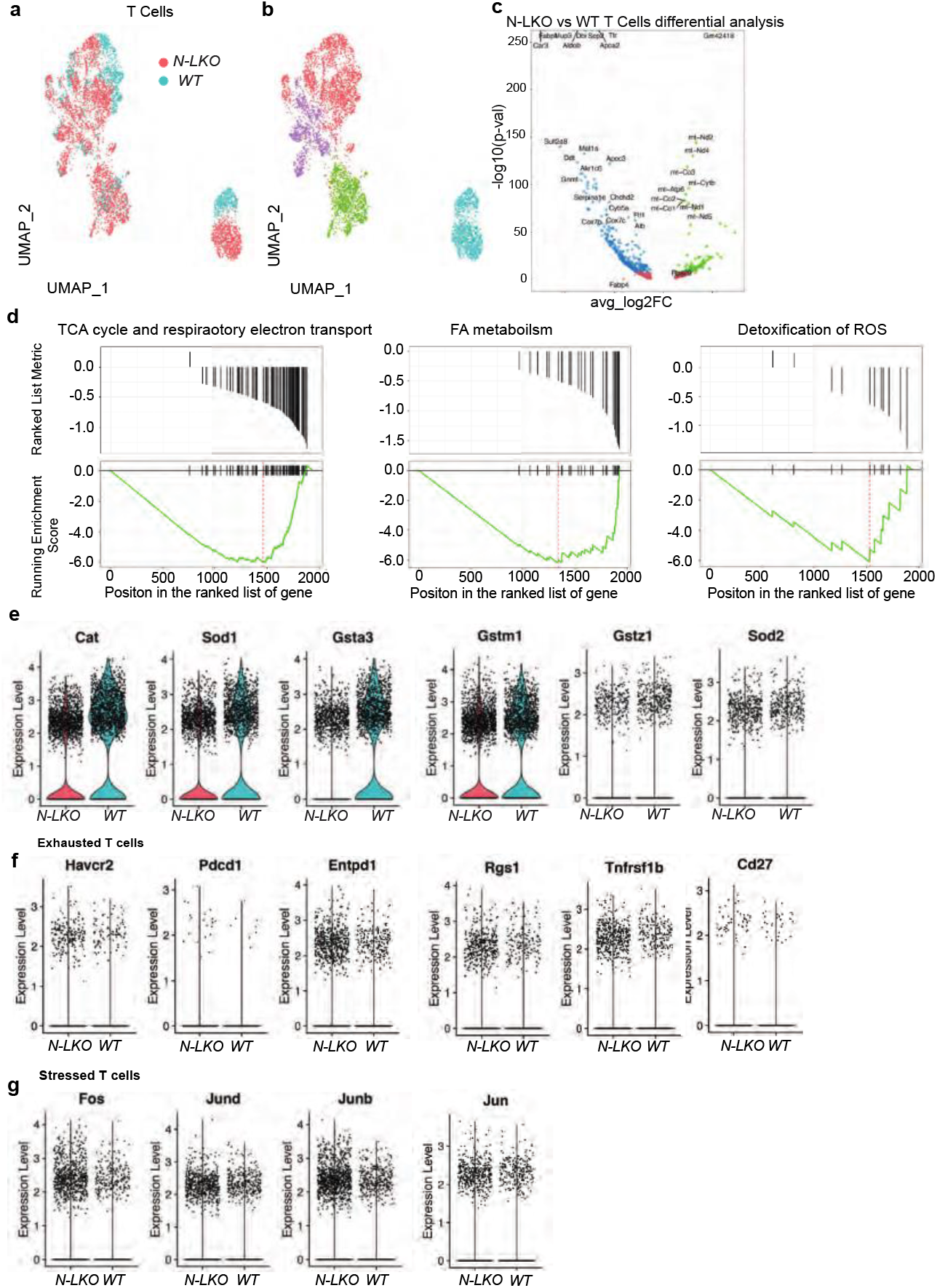
Hepatic loss of NgBR impacts on T cell profile. The single-cell RNA sequencing analysis of liver cells from WT and N-LKO mice fed a WD revealed important differences in the gene expression patterns of T cells. **a-b,** The UMAP plot displays four distinct subclusters of T cells from both groups, colored according to genotype. The color represents the subcluster or genotype. **c,** The Volcano plot illustrates the differential expression of genes in upregulated and downregulated T cells in N-LKO mice compared to WT mice. **d,** GSEA plot indicates the downregulation of genes involved in mitochondrial respiratory function, fatty acid metabolism, and detoxification of ROS in T cells of N-LKO mice compared to WT mice. **e,** The violin plots demonstrate that T cells from N-LKO mice had significantly downregulated antioxidant genes compared to those from WT mice. **f-g,** The Violin plots show a significant upregulation of genes associated with T cell exhaustion and stress in hepatic T cells of N-LKO mice compared to WT mice.

**Extended Data Fig. 15.**
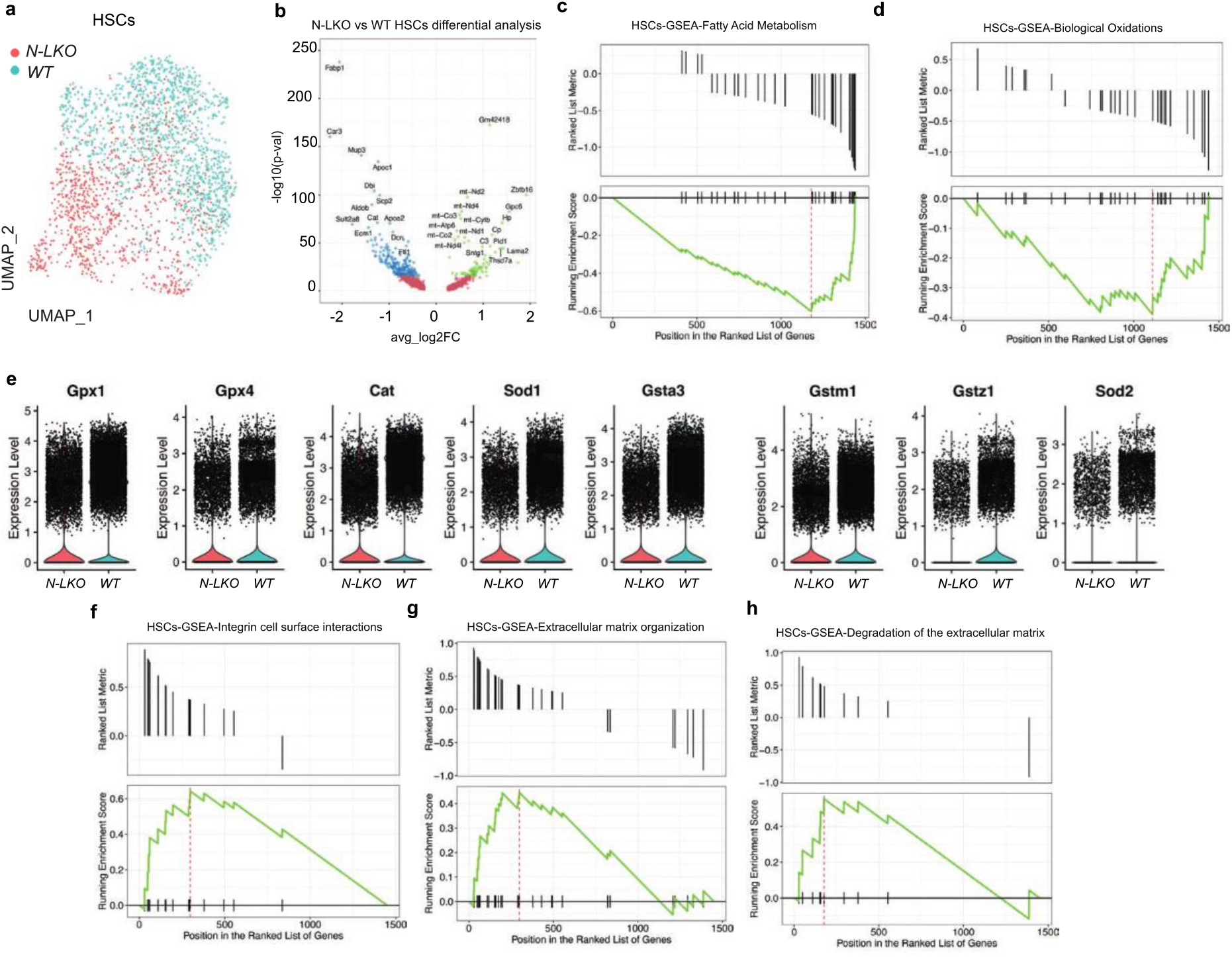
Loss of NgBR in hepatocytes triggers the activation of hepatic stellate cell gene profile. Impact of hepatic loss of NgBR on the hepatic stellate cell gene profile, as analyzed through single-cell RNA sequencing of liver cells from WT and N-LKO mice fed a Western diet (WD). **a,** The UMAP plot illustrates distinctive subclusters of hepatic stellate cells in both groups, which are distinguished by color. **b,** The volcano plots illustrate the differential expression of genes, demonstrating significant upregulation and downregulation of hepatic stellate cells in N-LKO mice compared to WT mice. **c-d,** Gene Set Enrichment Analysis (GSEA) plots that show the downregulation of HSC genes involved in lipid oxidation in N-LKO compared to WT mice **e,** The violin plots visually depicted the significant downregulation of expression levels of HSC genes involved in the oxidative stress response in N-LKO mice relative to WT mice. **f-h,** GSEA plots that show upregulation of HSC genes on extracellular cell matrix (ECM) remodeling

**Extended Data Fig. 16.**
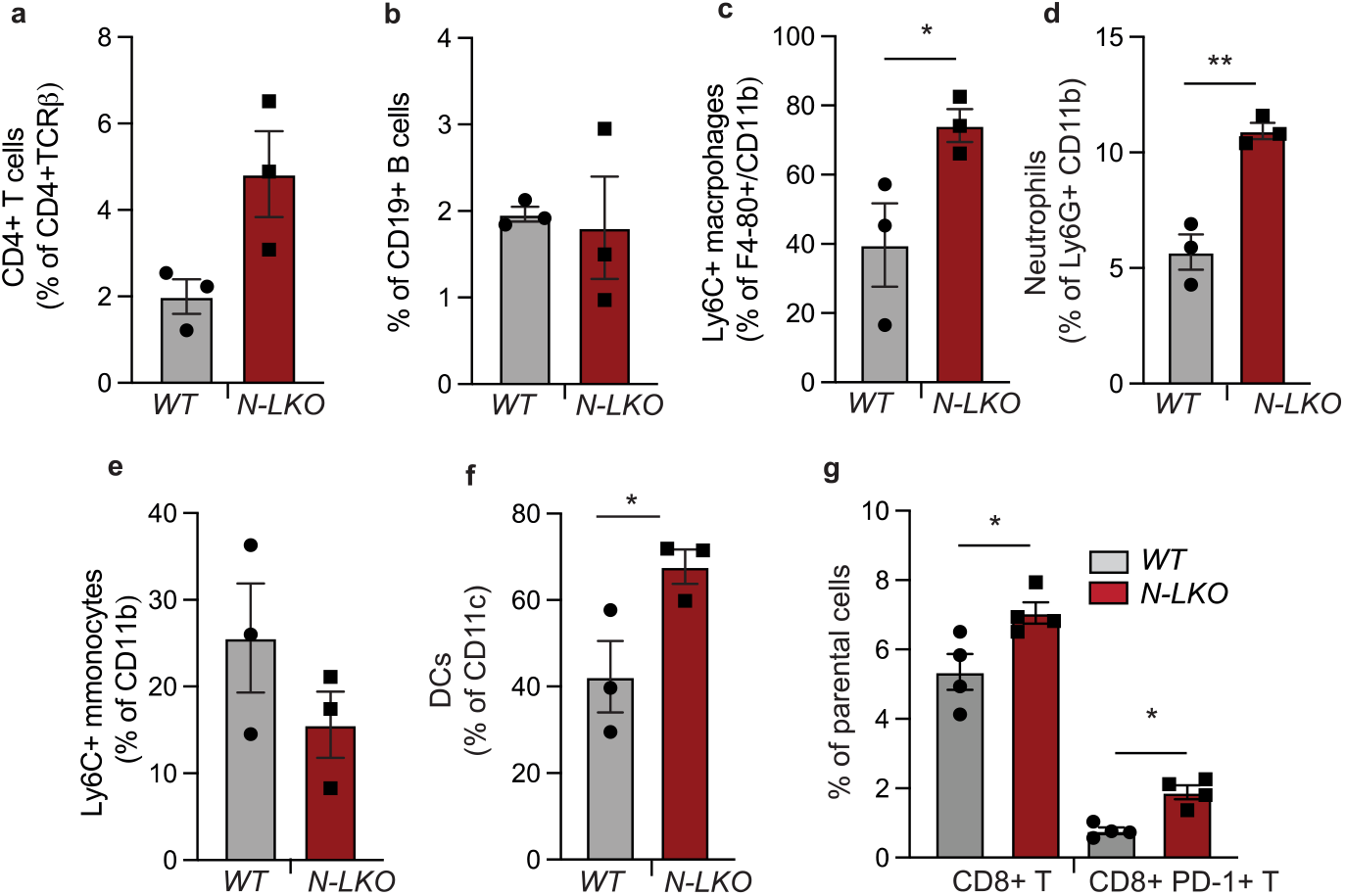
Lack of NgBR in the liver enhances hepatic immune infiltration. **a-g,** Hepatic lymphoid cells including CD4+ T cells and CD19+ B cells and myeloid cells including macrophages, neutrophil cells, monocytes and dendritic cells (DCs) and immunosuppressive CD8+PD1+ T cells isolated from WT and N-LKO mice fed an HFD for 16 weeks assessed by flow cytometry. Two-sided; \**P* < 0.05; \*\**P* < 0.01; comparing N-LKO with WT mice using an unpaired Welch’s t-test.

**Extended Data Fig. 17.**
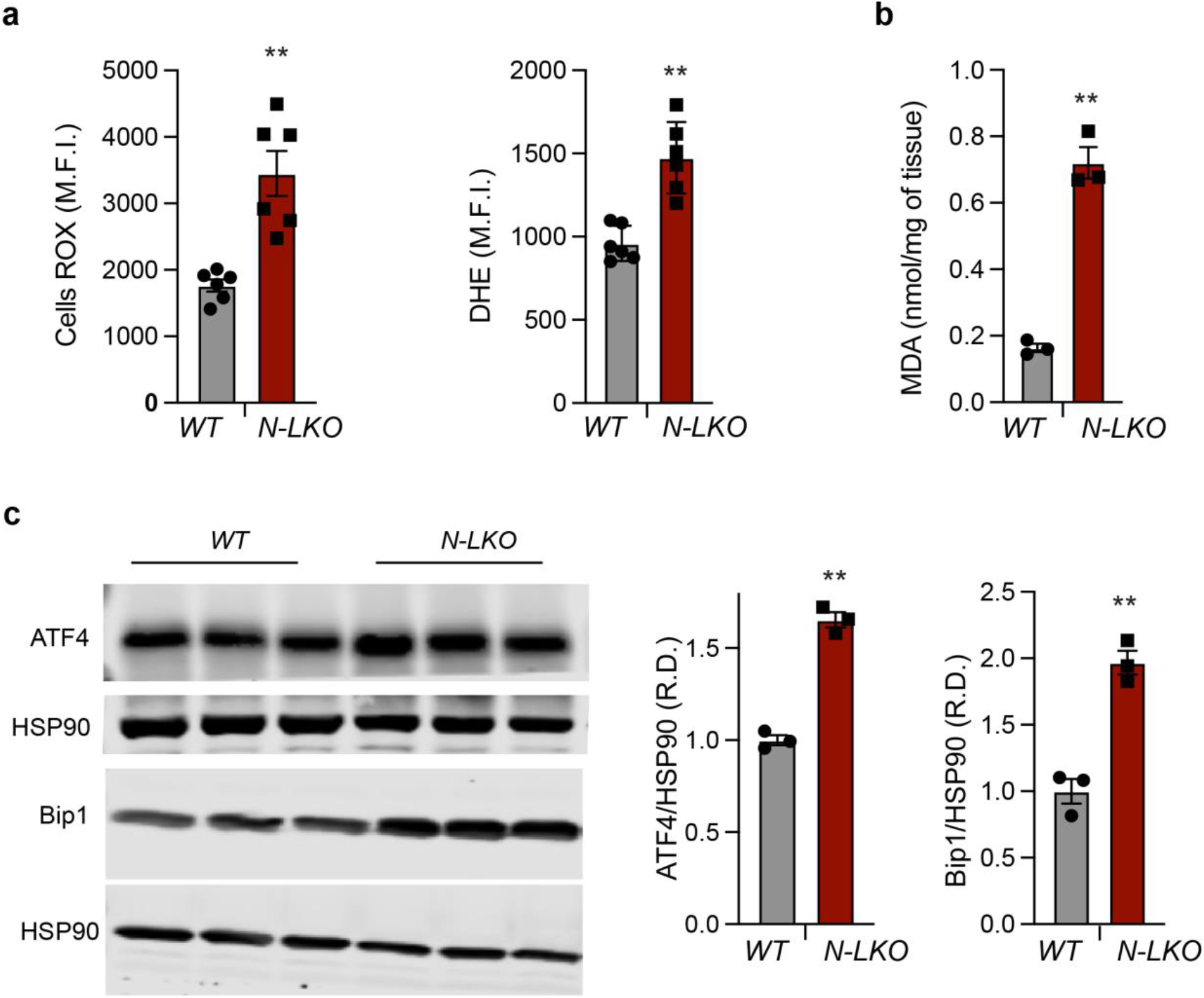
absence of NgBR in the liver induces oxidative and ER stress. **a,** Analysis of cellular ROS production in the primary hepatocytes isolated from WT, N-LKO fed a HFD (n=6). **b,** Membrane lipid peroxidation determined via MDA assay in the liver isolated from WT and N-LKO fed HFD (n=3). **c,** Representative immunoblot blot and densiometric analysis of an ER key stress response proteins ATF4 and Bip1 and housekeeping standard HSP90 in the liver isolated from WT and N-LKO fed HFD (n=3). Two-sided \*\**P* < 0.01; comparing N-LKO with WT mice using an unpaired Welch’s t-test.

